# *Legionella pneumophila* modulates host cytoskeleton by an effector of transglutaminase activity

**DOI:** 10.1101/2022.11.15.516567

**Authors:** Yan Liu, Yao Liu, Zhao-Qing Luo

## Abstract

The bacterial pathogen *Legionella pneumophila* delivers more than 330 effector proteins into host cells through its Dot/Icm type IV secretion system (T4SS) to facilitate its intracellular replication. A number of these effectors modulate organelle trafficking pathways to create a membrane-bound niche called the *Legionella* containing vacuole (LCV). In this study, we found that *L. pneumophila* induces F-actin accumulation in host cell cortex by its Dot/Icm substrate RavJ (Lpg0944). RavJ harbors an C_101_H_138_D_170_ motif associated with human tissue transglutaminases (TGs). We showed that RavJ catalyzes a covalent linkage between actin and the Motin family proteins Angiomotin (AMOT) and Angiomotin-like 1 (AMOTL1), proteins known to regulate tube formation and cell migration. Further study revealed that RavJ-induced crosslink between actin and AMOT occurs on its Gln_354_ residue. Crosslink between actin and AMOT significantly reduces the binding between actin and its binding partner cofilin, suggesting that RavJ inhibits actin depolymerization. We also demonstrated that the metaeffector LegL1 directly interacts with RavJ to antagonize its transglutaminase activity, leading to reduced crosslink between actin and Motin proteins. Our results reveal a novel mechanism of modulating the host actin cytoskeleton by *L. pneumophila*.

## Introduction

*Legionella pneumophila* is a Gram-negative intracellular pathogen that causes Legionnaires’ disease in humans[1]. Successful colonization by this bacteria requires its ability to manipulate such diverse processes of host cells as membrane trafficking, immunity, protein translation, autophagy, gene expression, and cytoskeleton structure[2, 3]. Upon entry into host cells, *L. pneumophila* promotes the biogenesis of a phagosome structure called the *Legionella*-containing vacuole (LCV) that supports its intracellular replication[4]. The virulence of *L. pneumophila* is correlated with its ability to survive and replicate in the LCV[4]. Biogenesis of the LCV requires the Dot/Icm system that transports over 330 protein substrates into the host cell[5, 6]. The activity of these effectors is essential for the development and maintenance of this replicative niche[4].

The actin cytoskeleton is involved in many essential cellular events such as mitosis, cell migration, control of epithelial barrier function, and adherence of immune cells[7]. Given the essential roles of the actin cytoskeleton, it is not surprising that this network is a common target for bacterial virulence factors[8]. *Yersinia* blocks macrophage phagocytosis by interfering with host Rho GTPase and the actin cytoskeleton dynamics[9]. *Salmonella enterica* Typhimurium delivers a subset of bacterial effector proteins into the eukaryotic host cell to modulate the host cell actin cytoskeleton, facilitating its own internalization into non-phagocytic cells[10]. *L. pneumophila* has also acquired the ability to modulate the actin cytoskeleton by its Dot/Icm substrates. For example, RavK is a protease that disrupts host cytoskeletal structure by cleaving actin[11]; LegK2 targets the actin nucleator ARP2/3 complex by phosphorylating its components ARPC1B and ARP3, leading to the global actin cytoskeleton remodeling in cells[12]; Ceg14 affects actin distribution and inhibits actin polymerization by a yet unknown mechanism[13]; VipA interferes with organelle trafficking by acting as a nucleator for actin polymerization[14].

Post-translational modifications (PTMs) of host proteins involved in important cellular processes is a commonly used mechanism used by bacterial pathogens to counteract host defense[15]. PTMs often is executed by virulence factors that display diverse biochemical activities. A number of unique PTMs have been found to be imposed by Dot/Icm effectors, including phosphorylcholination[16, 17], AMPylation[18], phosphorylation[19], ADP-ribosylation[20–22], ubiquitination[20, 23], and transglutamination[23]. Among these, transglutamination is induced by transglutaminases (TGs) that primarily catalyze the formation of an isopeptide bond between the γ-carboxamide group of a glutamine residue from one protein and the ε-amino group of a lysine residue of another protein with the release of an ammonia[24]. This modification has been identified as an important PTM that attacks a wide spectrum of host functions to benefit the pathogens. For example, the HopX (AvrPphE) family of *Pseudomonas syringae* Type III effectors are composed of a conserved putative cysteine-base catalytic triad resemble of the transglutaminase family that is required for the generation of a cell-death response in specific *Arabidopsis* ecotypes[25]. The type III effector VopC promotes *Vibrio* invasion by activating Rac and CDC42 via its transglutaminase activity[26]. Modification by transglutamination by *L. pneumophila* effectors has recently emerged as an important virulence factor of this bacterium. The *L. pneumophila* effector MavC has been characterized as a transglutaminase that catalyzes the formation of a covalent linkage between ubiquitin and UBE2N, leading to the inhibition of NF-κB signaling in the initial phase of bacterial infection[23].

Here, we showed that the Dot/Icm substrate RavJ (Lpg0944) catalyzes crosslink between actin and members of the Motin family, AMOT and AMOTL1, via its transglutaminase activity, leading to accumulation of actin polymers in mammalian cells. We also showed that the transglutaminase activity of RavJ is regulated by another Dot/Icm substrate LegL1(Lpg0945), which directly binds to RavJ and inhibits the enzymatic activity of RavJ.

## Results

### RavJ is a transglutaminase that induces actin accumulation in mammalian cells

RavJ was originally identified in a study aiming to analyze the mechanism of metaeffector activity in *L. pneumophila*[27]. Structural analysis reveals that RavJ harbors a C-H-D (C_101_H_138_D_170_) motif associated with members of tissue transglutaminases (TGs)[27]. In a screening to identify Dot/Icm substrates capable of regulating the actin cytoskeleton, we transfected HEK293T cells with a GFP fusion library of Dot/Icm substrates and found that RavJ causes rearrangements of the actin cytoskeleton in mammalian cells (**Fig 1**). To determine the role of the putative catalytic motif potentially involved in TG activity in the function of RavJ, we introduced mutations in C_101_, H_138_, D_170_, respectively, and found that each of these mutations completely abolished the ability of RavJ to induce actin cytoskeleton rearrangements (**Fig 1**). These results indicate that overexpression of RavJ triggers the formation of actin filaments in the cell cortex by a mechanism that requires its C_101_H_138_D_170_ motif.

**Fig 1.**
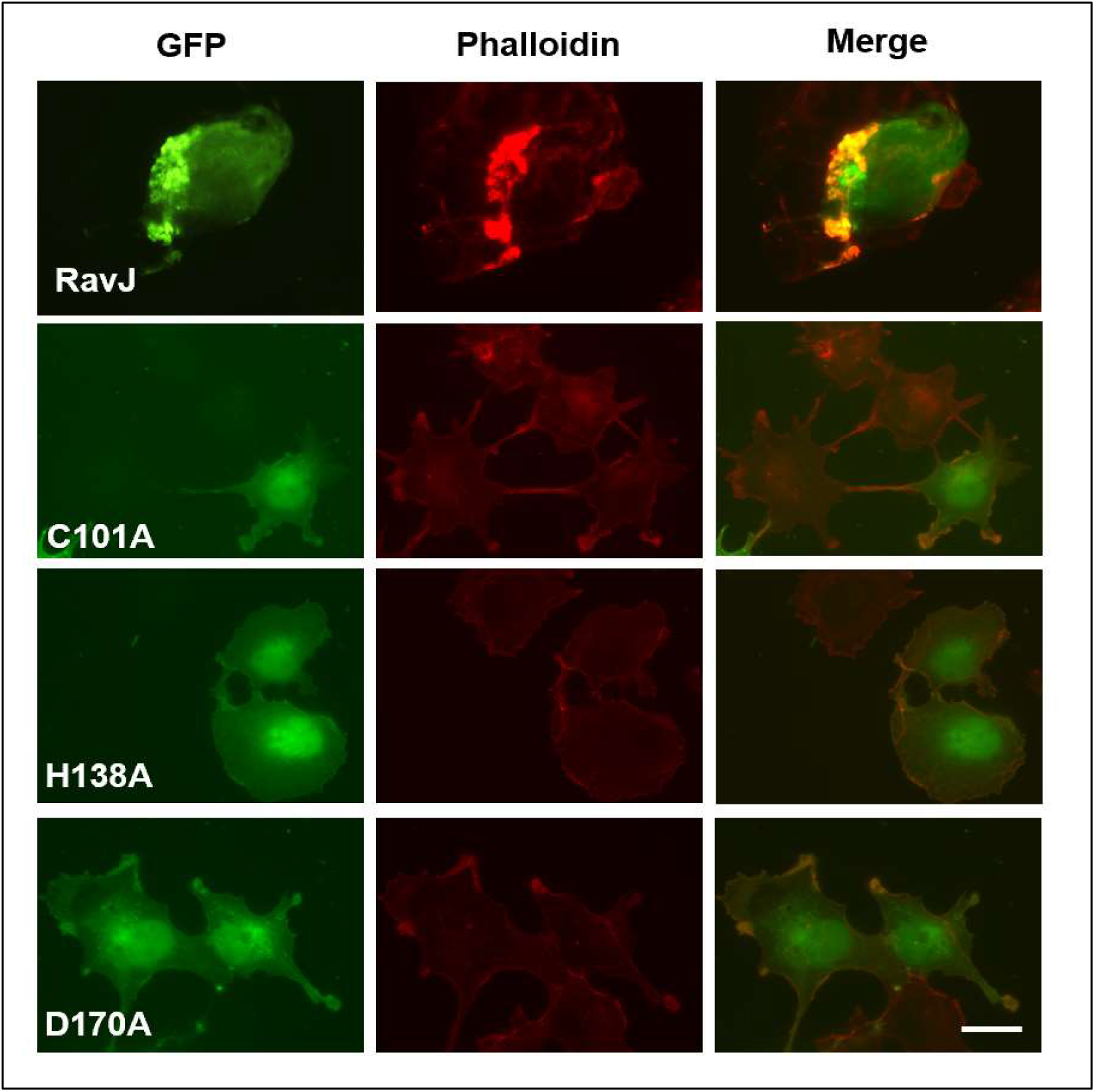
Ectopic expression of RavJ causes rearrangement of actin cytoskeleton. HEK293T cells were transfected with the indicated constructs and then subjected to immunofluorescence microscopic analysis. F-actin was stained by phalloidin conjugated with Texas-Red. Bar, 10 μm. Note that wild-type RavJ induces actin accumulation in the cell cortex and this phenotype is dependent on the predicted TG enzymatic motif.

### *ravJ* is dispensable for intracellular growth of *L. pneumophila*

To examine the role of *ravJ* in *L. pneumophila* virulence, we first determined the level of RavJ at different growth phases throughout its growth cycle in broth. RavJ was detectable in all growth phases (optical density at 600 nm (OD_600_) of 0.5-3.5) but became highly expressed in the lag phase (OD_600_=0.5-0.7) (**Fig 2A**) after saturated cultures were diluted into fresh medium, suggesting that RavJ functions in the initial phase of infection.

**Fig 2.**
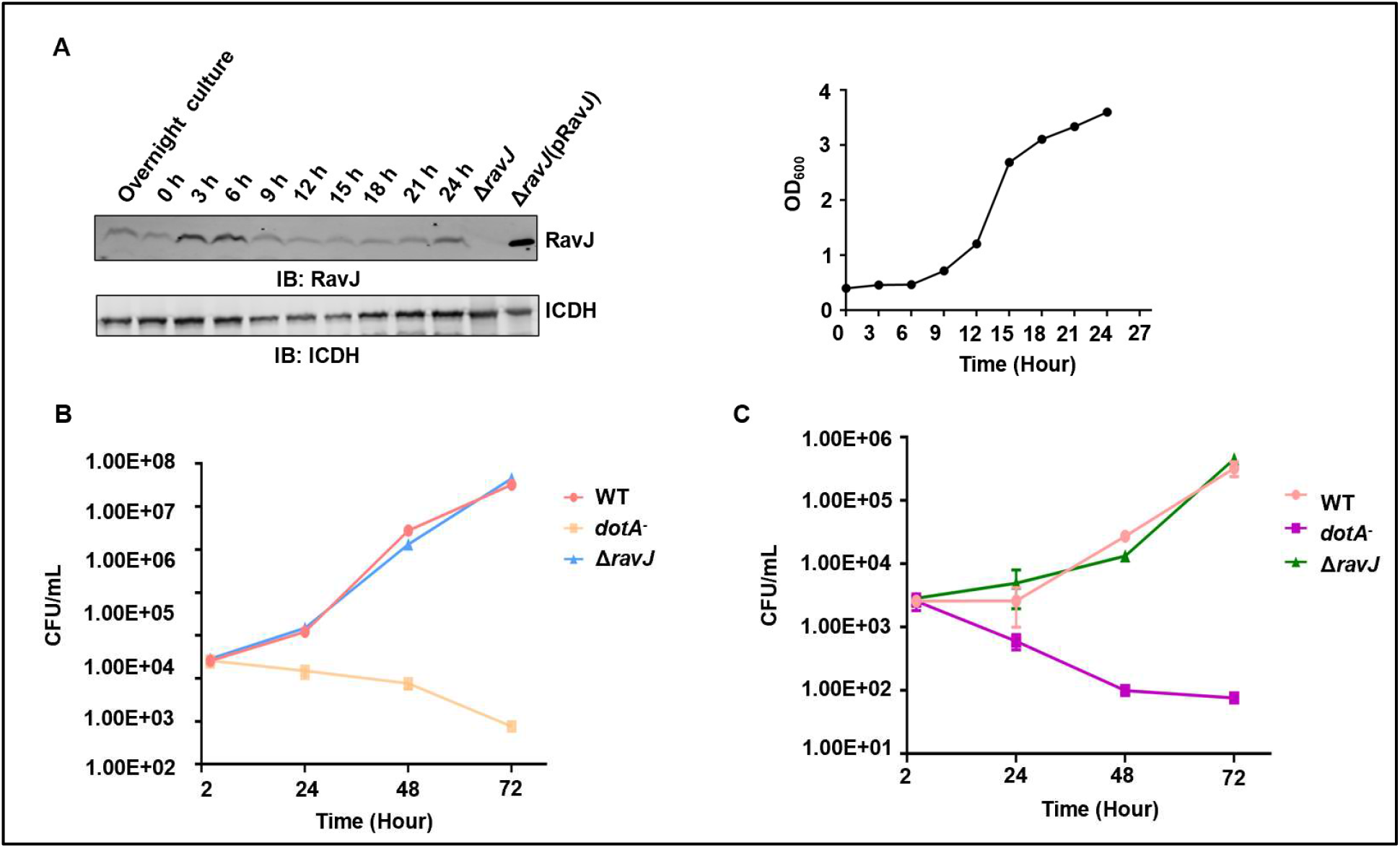
*ravJ* does not influence intracellular growth of *L. pneumophila*. **A**. The growth of *L. pneumophila* in AYE broth (right) and the expression of RavJ in bacteria grown in broth (left). Cultures grown to stationary phase were diluted 1:20 into fresh medium and the growth of bacteria was monitored by measuring OD_600_ at the indicated time points. **B-C**. The Δ*ravJ* strain grew at rates indistinguishable from that of the wild-type strain in commonly used tissue culture hosts. Bone marrow-derived mouse macrophages (BMDMs) **(B)** or *Dictyostelium discoideum* **(C)** were infected with relevant *L. pneumophila* strains at the indicated time points, cells were treated with 0.02% saponin for half an hour and the bacteria number was determined by enumerating colony-forming unit (CFU) of appropriately diluted lysates obtained by saponin. Errors bars represent ±SEM; n=3;

We also determined the role of *ravJ* in *L. pneumophila* virulence by examining intracellular bacterial replication of relevant bacterial strains. The mutant Lp02ΔravJ grew at rates indistinguishable from that of the wild-type strain (**Fig 2B-C**), indicating that, similar to most Dot/Icm substrates, RavJ is not required for proficient intracellular bacterial replication in commonly used tissue culture hosts.

### Ectopically expressed RavJ crosslinks with actin

TGs function to catalyze homo- or heterologous protein crosslink by a transglutamination reaction[24]. The key to understand the mechanism of how RavJ induces actin rearrangement is to identify its target proteins. To achieve this goal, we expressed Flag-tagged RavJ or the RavJ_C101A_, RavJ_H138A_, RavJ_D170A_ mutants in HEK293T cells. Cell lysates were subjected to immunoprecipitation with beads coated with the Flag antibody. Detection of the precipitated proteins by immunoblotting with the Flag antibody revealed that a portion of wild-type RavJ but not the mutants migrated as higher molecular weight (MW) forms (**Fig 3A**), suggesting it is posttranslationally modified. To identify the modification associated with the upshifted protein, we expressed double-tagged Flag-RavJ-His_6_ in HEK293T cells and used a two-step sequential purification procedure to obtain upshifted RavJ (**Fig 3B**). Mass spectrometric analysis of proteins in the upshifted band identified abundant actin and RavJ, suggesting that the upshifted band is a product generated by protein conjugation between these two proteins.

**Fig 3.**
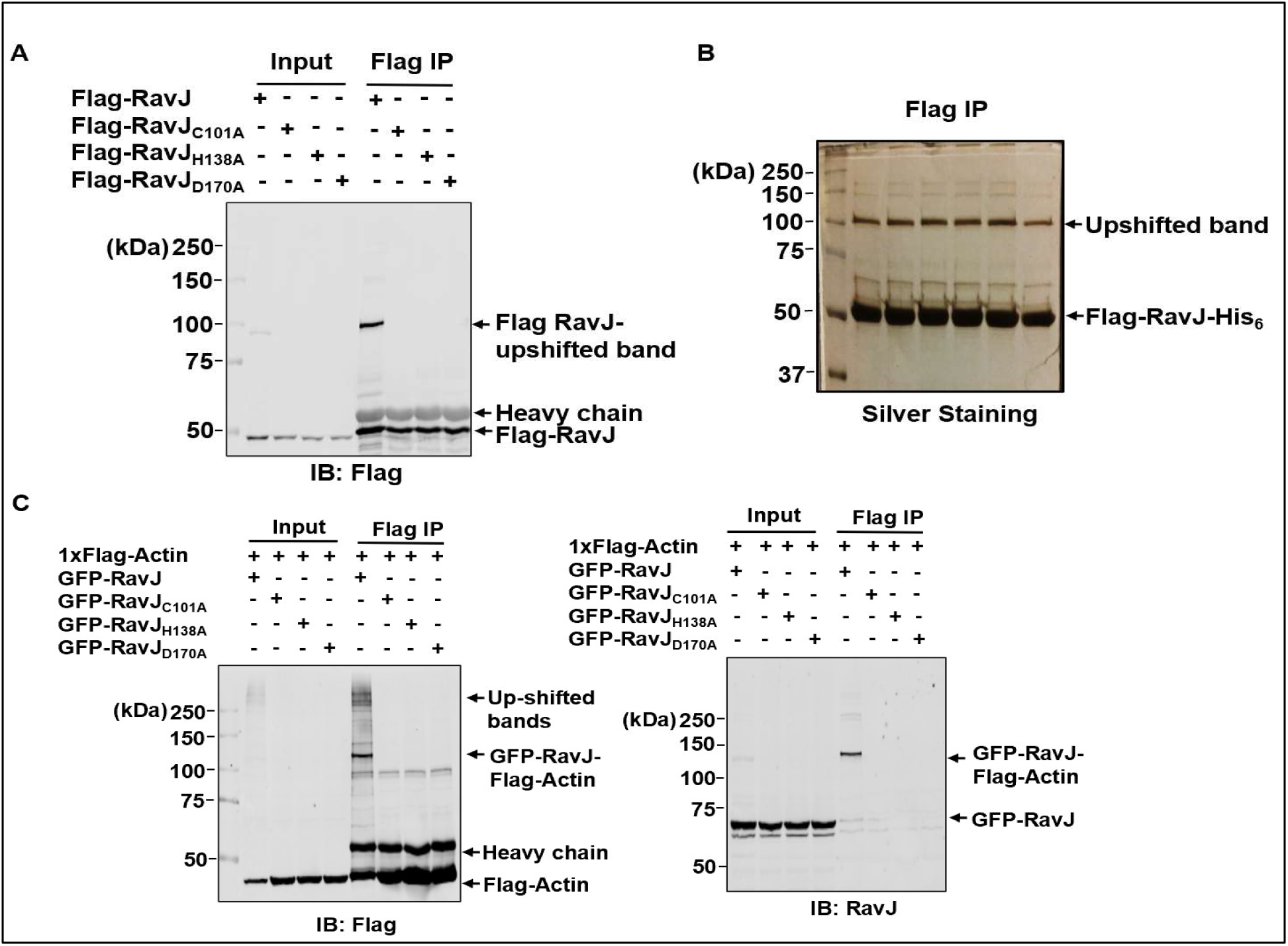
RavJ induces a molecular weight shift in actin by a putative transglutaminase activity. **A**. A portion of RavJ migrated as a higher molecular weight form when expressed in mammalian cells. HEK293T cells were transfected to express Flag-RavJ, RavJ_C101A_, RavJ_H138A_, or the RavJ_D170A_ mutants. Total cell lysates were immunoprecipitated (IP) with beads coated with the Flag antibody. Products were resolved by SDS-PAGE and probed with the Flag antibody. Note that RavJ displayed as an upshifted band on the blot. **B**. Tandem purification of the Flag-RavJ-His_6_ upshifted band. HEK293T cells were transfected to express Flag-RavJ-His_6_. Cell lysates were subjected to IP with beads coated with the Flag antibody. Proteins were then eluted from the beads with 3xFLAG peptide and incubated with Ni^2+^-NTA beads. Products were separated by SDS-PAGE and detected by silver staining. **C**. RavJ forms a covalent bond with actin in mammalian cells. HEK293T cells expressing the indicated proteins were subjected to IP with beads coated with the Flag antibody and separated by SDS-PAGE. Samples were detected by Flag-specific (left) and RavJ-specific (right) antibody, respectively.

To test whether the upshifted band is a crosslink product between actin and RavJ, we coexpressed Flag-tagged actin and GFP fusion of RavJ or its mutants in HEK293T cells. Cell lysates were subjected to immunoprecipitation using agarose beads coated with the Flag-specific antibody. We found that GFP-RavJ indeed was linked with actin (**Fig 3C**). To identify the chemical linkage between actin and RavJ, the protein band corresponding to the RavJ upshifted band was excised, digested with trypsin, and analyzed by mass spectrometry (MS). TGs catalyze protein crosslinking by forming an isopeptide bridge between the lysine (Lys) donor residue of one protein and the acceptor glutamine (Gln) residue of another[24]. Thus, we hypothesized that a lysine residue and a glutamine residue are involved in the crosslink between RavJ and actin. In the MS analysis, around 80% of the peptides in actin were recovered from the tryptic digestion, indicating Lys_84_, Lys_118_, Lys_213_, Lys_359_, Gln_121_, Gln_137_, Gln_353_, Gln_354_ in actin as potential lysine donor residues or glutamine acceptor residues (**Fig 4A**). Among the individual mutation of these residues, only Q354A in actin abolished the ability of actin to crosslink with RavJ (**Figs 4B and S1A-B**).

**Fig 4.**
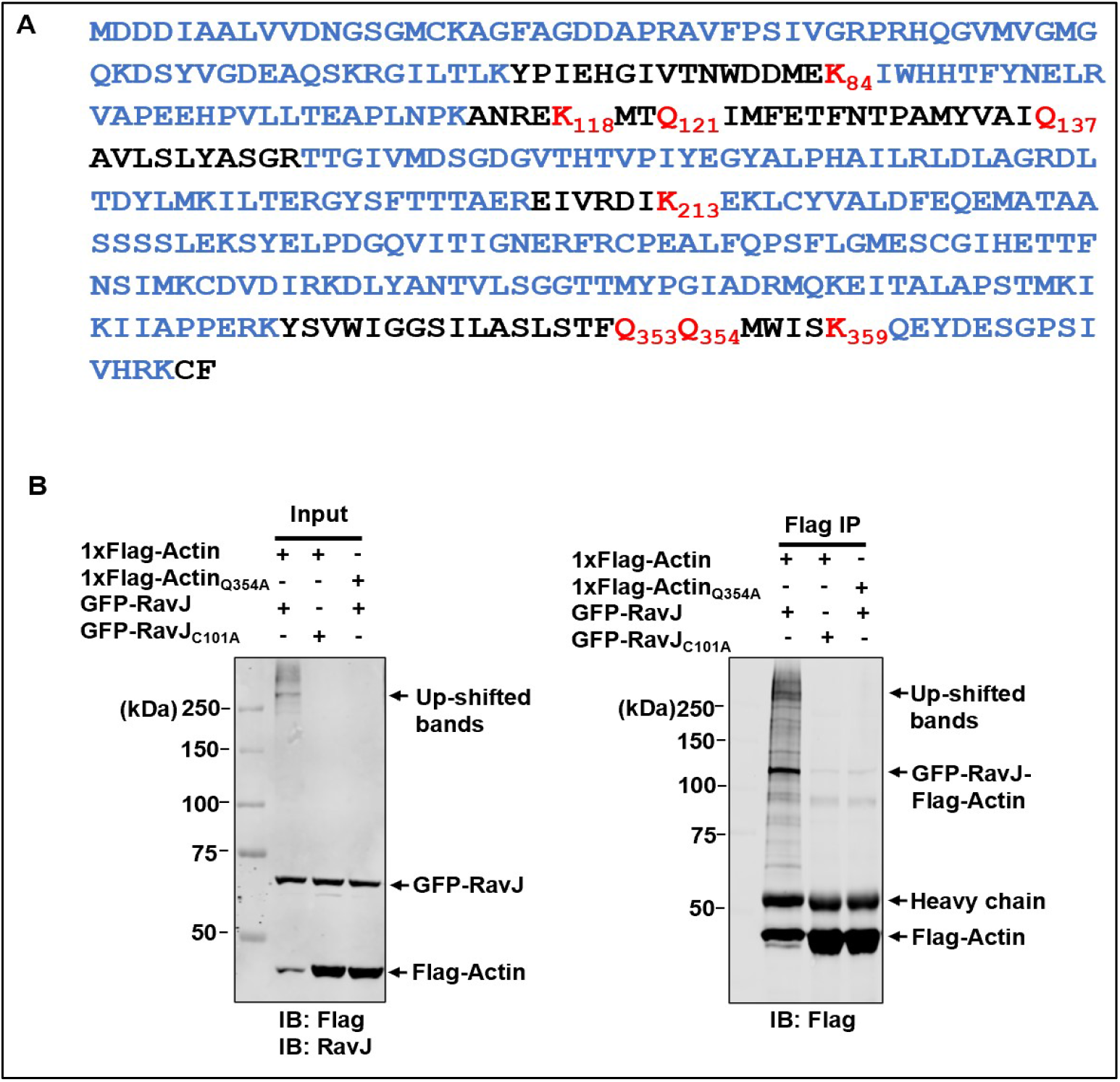
Actin is crosslinked to RavJ through its Gln_354_ residue. **A.** MS analysis of the Flag-RavJ-His_6_ upshifted band. Coverage of the amino acids in actin are highlighted in blue. Note that Lys_84_, Lys_118_, Lys_213_, Lys_359_, Gln_121_, Gln_137_, Gln_353_, Gln_354_ were not recovered from the MS, indicating them as potential linkage residues. **B.** The formation of Actin-RavJ was detected by immunoblotting. Note that the Actin Gln_354_Ala mutant has largely lost the ability to be modified (the 3^rd^ lane).

### RavJ induces crosslink between actin and members of the Motin family protein

Transglutaminases normally catalyze crosslink between two proteins, and in the absence of the receptor substrate, these enzymes induce crosslink between itself and the available donor substrate. Our analysis of the precipitated products obtained by RavJ identified a number of proteins (**Fig 3C**), one or more of which could be potentially the second substrate that crosslinks with actin in the reaction induced by RavJ. To identify such proteins, we purified the crosslink products using a tandem purification method from cells transfected to express Flag-HA-actin and GFP-RavJ. Samples similarly transfected to express the catalytically inactive RavJ_C101A_ mutant were used as controls. Twenty-four hours after transfection, cell lysates were subjected to IP with beads coated with the Flag antibody. Proteins eluted with 3XFLAG peptide were further purified by IP with the HA antibody. Samples separated by SDS-PAGE were detected by immunoblotting with the appropriate antibodies. In samples transfected to express GFP-RavJ, several upshifted bands were detected with the anti-Flag antibody (**S2A Fig**). The gels containing upshifted proteins were excised, digested with trypsin, and analyzed by mass spectrometry, which allowed us to obtain a list of proteins that potentially crosslink with actin (**Table 1**). Two of Wiskott-Aldrich syndrome and SCAR homolog (WASH) complex components, WASHC4 and WASHC5, and one protein from the Motin family, Angiomotin (AMOT) were among the proteins identified with high confidence. We considered these proteins as potential substrates of RavJ because of their relevance to the actin cytoskeleton network. Further analysis showed that RavJ did not detectably induce crosslink between actin and WASHC4 or WASHC5 (**S2B-2C Fig**). Importantly, we found that AMOT is able to form a conjugate with actin when coexpressed with RavJ (**Fig 5B**), suggesting that AMOT is the second substrate of the crosslinking reaction catalyzed by RavJ.

**Fig 5.**
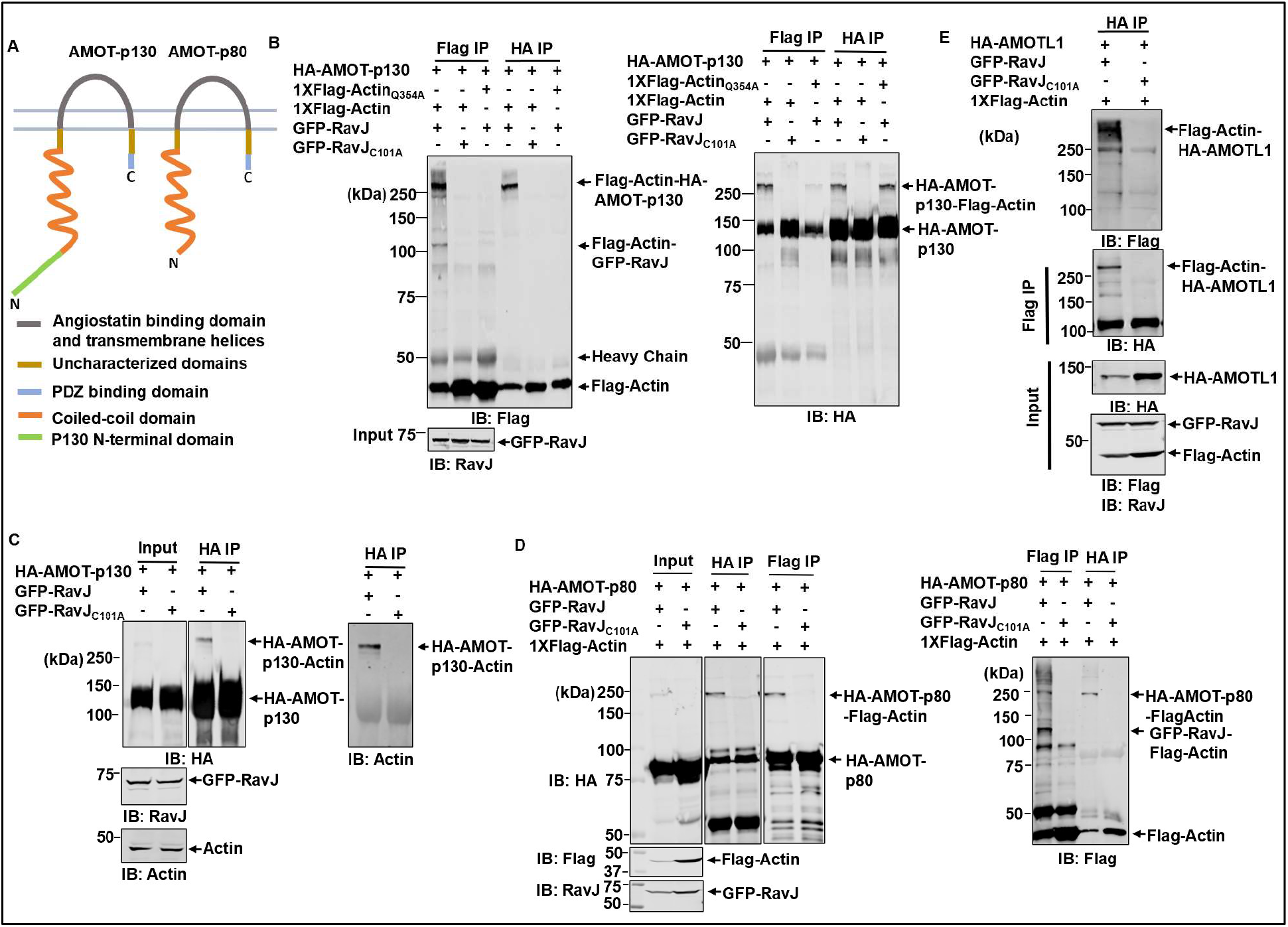
RavJ catalyzes crosslinks between actin and members of the Motin protein family. **A**. A schematic view of the two isoforms of AMOT produced by alternative splicing of the *AMOT* mRNA. The two isoforms, AMOT-p130 and AMOT-p80, are characterized by a conserved coiled-coil domain, a C-terminal PDZ binding domain, and an angiostatin binding domain. The AMOT-p130 isoform harbors a unique N-terminal domain. **B**. Actin conjugates with AMOT-p130 in the presence of RavJ. HEK293T cells co-transfected with the indicated proteins were lysed and subjected to IP with beads coated with the Flag antibody or the HA antibody. Samples resolved by SDS-PAGE were probed with a Flag-specific antibody or an HA-specific antibody. **C**. RavJ induces crosslink between HA-AMOT and endogenous actin. Cells expressing HA-AMOT-p130 and GFP-RavJ or the GFP-RavJ_C101A_ mutant were lysed and immunoprecipitated with beads coated with the HA antibody. Crosslink products were probed by immunoblotting using either Actin or Flag antibody. **D**. Actin forms a conjugate with AMOT-p80 in HEK293T cells in the presence of RavJ. Cells transfected to express the indicated proteins were lysed and subjected to IP with beads coated with Flag or HA antibody, followed by immunoblotting analysis. The proteins were detected by antibodies specific to a Flag, RavJ or HA. **E**. RavJ catalyzes the crosslink between AMOTL1 and actin. Cells transfected to express the indicated proteins were lysed and subjected to reciprocal IP, followed by SDS-PAGE analysis, and was probed with the indicated antibodies.

**Table 1:** Top hits from MS analysis of the tandem purification.

| Protein Names | Ratio (RavJ/RavJ<sub>C101A</sub>) | MS/MS count RavJ<sub>C101A</sub> | MS/MS count RavJ | Mol. weight [kDa] |
| --- | --- | --- | --- | --- |
| Actin | 29.68 | 25 | 742 | 42 |
| Splicing factor, proline- and glutamine-rich | 19.5 | 2 | 39 | 76 |
| Kinesin-like protein KIF11 | 17.5 | 6 | 105 | 119 |
| Adenylyl cyclase-associated protein 1 | 7.78 | 14 | 109 | 51 |
| Cytoplasmic dynein 1 heavy chain 1 | ∞ | 0 | 155 | 532 |
| Eukaryotic translation initiation factor 4 gamma 1 | ∞ | 0 | 84 | 158 |
| Filamin-A | ∞ | 0 | 63 | 280 |
| WASH complex subunit strumpellin (WASHC5) | ∞ | 0 | 52 | 134.28 |
| Talin-1 | ∞ | 0 | 44 | 270 |
| Insulin receptor substrate 4 | ∞ | 0 | 42 | 133 |
| WASH complex subunit 7 (WASHC4) | ∞ | 0 | 35 | 136.4 |
| Afadin | ∞ | 0 | 22 | 210 |
| Eukaryotic translation initiation factor 4 gamma 2 | ∞ | 0 | 19 | 98 |
| General transcription factor II-I | ∞ | 0 | 18 | 108 |
| Vigilin | ∞ | 0 | 17 | 141 |
| Src substrate cortactin | ∞ | 0 | 16 | 57 |
| Angiomotin | ∞ | 0 | 5 | 130 |

To verify that AMOT is targeted by RavJ, we tested the crosslink between endogenous actin and AMOT. In cells transfected to express GFP-RavJ, crosslink products formed by endogenous actin and AMOT were detected (**Fig 5C**). We also observed crosslink between actin and AMOT-p80 (**Fig 5D**), an AMOT isoform arose from alternative splicing of the *AMOT* transcript (**Fig 5A**). In addition, we have tested Angiomotin-like 1 (AMOTL1), which is another member in the Motin family having significant homology with AMOT. Crosslink products of AMOTL1 and actin were detected in HEK293T cells expressing wild-type RavJ but not its catalytically inactive mutants (**Fig 5E**). Taken together, our results indicate that RavJ catalyzes the crosslink between actin and members of the Motin family.

The Motin family of proteins harbor several structural domains, including the N-terminus domain potentially involved in Yes-associated protein 1 (YAP1) binding[28], a conserved coiled-coil (CC) domain and the C-terminal PDZ-binding domain[29] (**Fig 6A**). To determine the crosslink site on AMOT, we constructed a number of HA-tagged AMOT truncation mutants (**Fig 6A**) and tested their ability to crosslink with actin in cells expressing RavJ. Our results showed that all the four truncations were able to crosslink with actin (**Fig 6B-C and 6F**). To identify the modification sites on each truncation, the corresponding crosslink products were subjected to MS analysis (**S3A-C Fig**). However, despite multiple attempts using different enzymes to digest the crosslink products, we were unable to detect the crosslink sites between the two proteins. The mass of the peptide without the crosslink is too big to extract the peptide from the gel and to sequence it. We then mutated the two lysine residues in the truncation contains the PDZ-binding domain (**Fig 6E**). Substitution mutant in Lys_1051_ was enough to abolish the crosslink product (**Fig 6E**). The same mutation in the AMOT_871-1084_ truncation also abolished the crosslink product (**Fig 6D**), suggesting that AMOT_871-1084_ crosslinks with actin through its Lys_1051_ residue.

**Fig 6.**
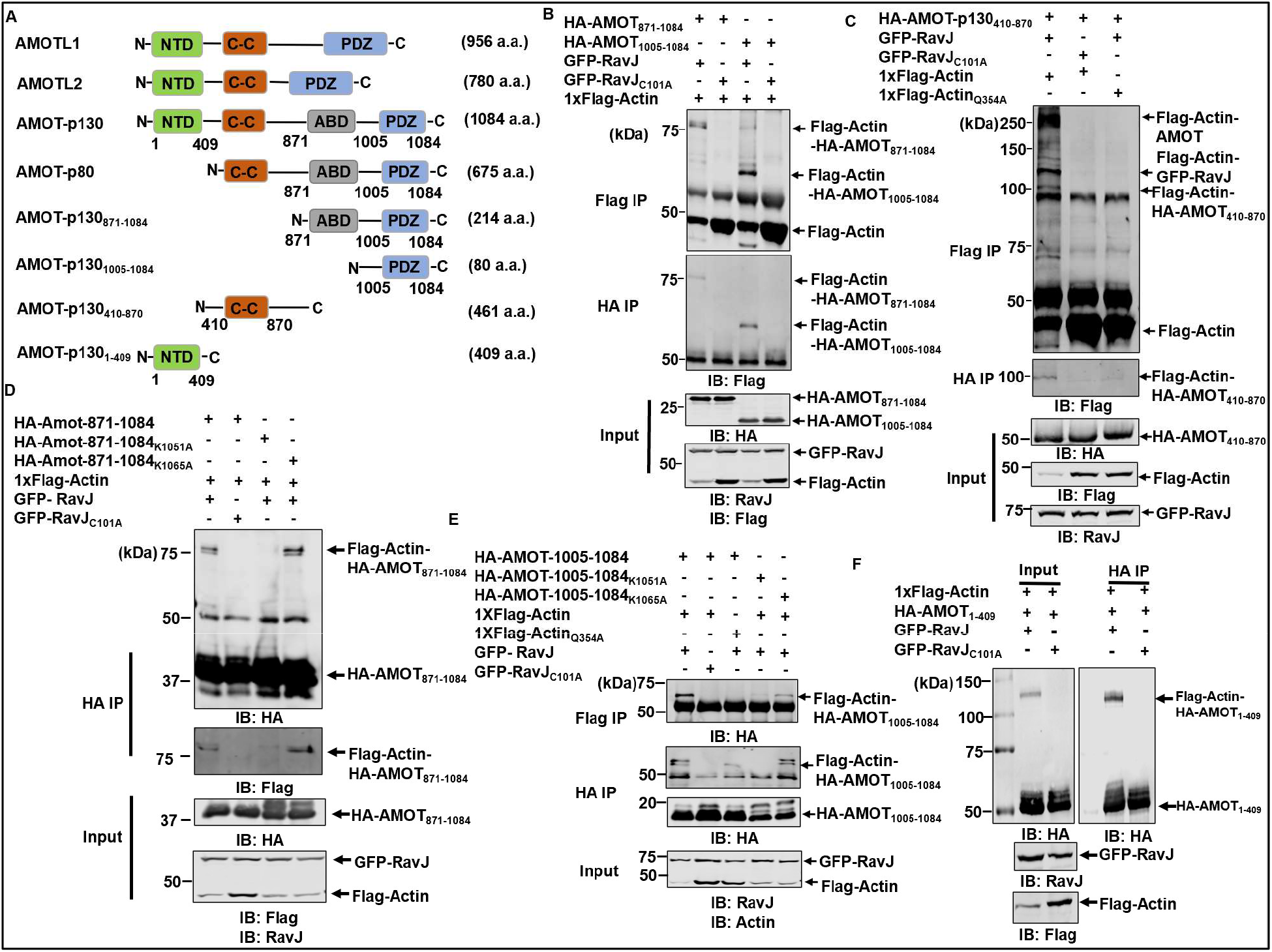
Determination of the modification sites on AMOT. HEK293T cells were transfected to express the indicated proteins. Cell lysates were subjected to reciprocal co-IP with Flag or HA antibody coated beads. Crosslink products were detected by immunoblotting with the indicated antibodies. **A**. Domain architecture of the Motin family of proteins and the AMOT-p130 truncations. The HA-tagged truncation mutants are made as indicated. **B**. The PDZ binding domain is enough to crosslink with actin. **C**. The AMOT coiled-coil truncation (AMOT_410-870_) crosslinks with actin in the presence of RavJ. **D**. The Lys_1051_Ala mutation in the AMOT_871-1084_ truncation abolished its ability to crosslink with actin. **E**. The Lys_1051_Ala mutation in the PDZ binding domain lost its ability to crosslink with actin. **F**. RavJ catalyzes the formation of a covalent bond between the N-terminal domain of AMOT-p130 and actin.

### RavJ catalyzes crosslink between purified actin and AMOT

We next examined whether crosslink between actin and AMOT occurs in cell-free reactions. HEK293T cellstransfected to express Flag-Actin, HA-AMOT, GFP-RavJ or GFP-RavJ_C101A_, respectively, were lysed, and the lysates were mixed and incubated at 37°C for 2 h. Beads coated with the Flag antibody were used to enrich Flag-Actin and products of crosslinking formed by actin and AMOT were detected by immunoblotting. Protein conjugate detectable by the HA-specific antibody was detected only in reactions that received lysates containing wild-type RavJ (**Fig 7A**), indicating that RavJ-induced crosslink between actin and AMOT occurs in a cell-free system.

**Fig 7.**
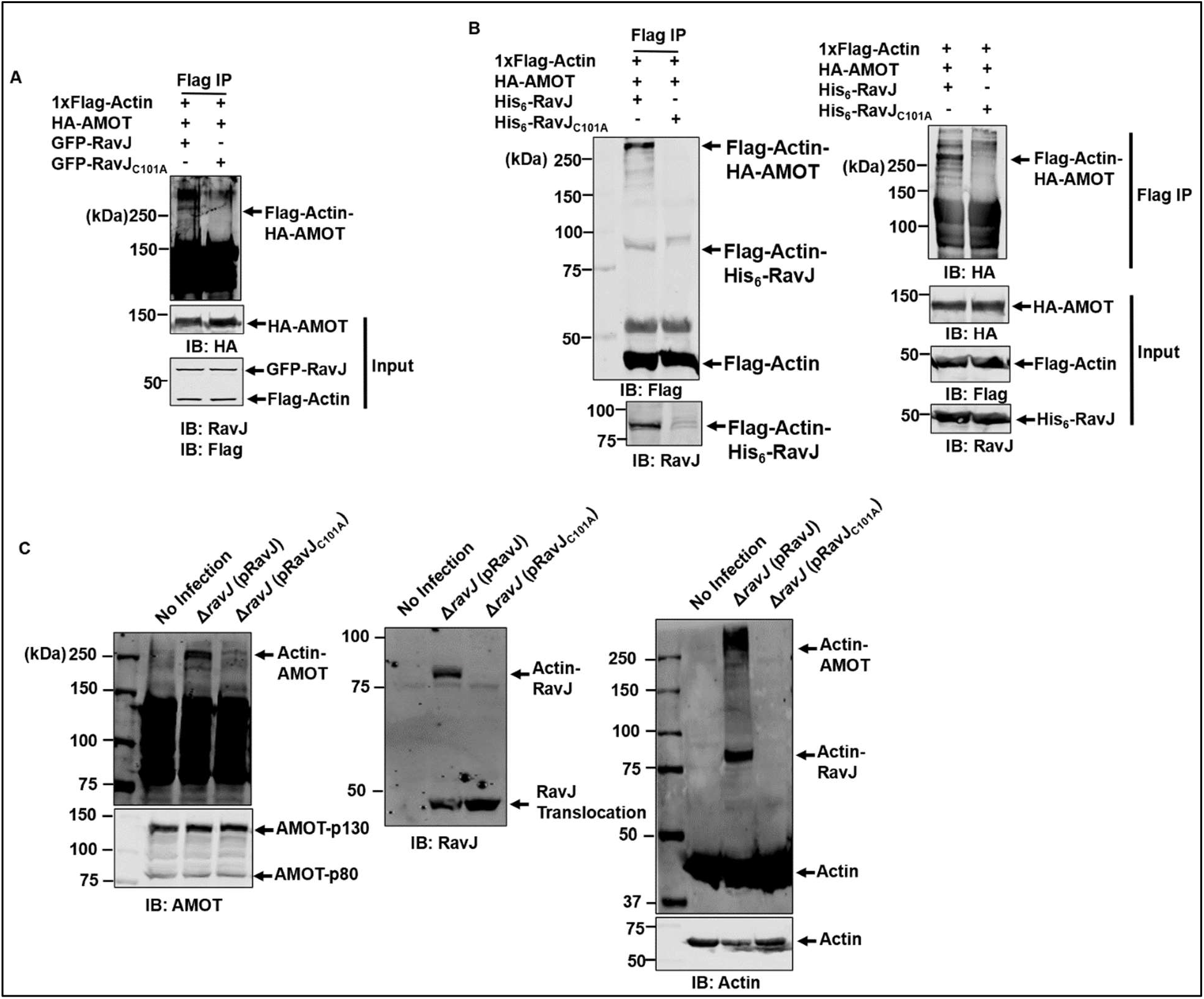
RavJ induces the formation of a protein conjugate by actin and AMOT. **A**. RavJ catalyzes crosslink between actin and AMOT in a cell-free system. Cells transfected to express Flag-Actin, HA-AMOT, GFP-RavJ, or GFP-RavJ_C101A_, respectively, were lysed with RIPA buffer without EDTA. Cell lysates were combined as indicated and incubated at 37°C for 2 h. Products were subjected to IP with beads coated with the Flag antibody, followed by immunoblotting with the indicated antibodies. **B**. RavJ induces crosslink between purified actin and AMOT *in vitro*. Reactions containing Flag-Actin, HA-AMOT, His_6_-RavJ or His_6_-RavJ_C101A_ were incubated at 37°C for 2 h. Samples were immunoprecipitated with beads coated with the Flag antibody. The crosslink products were detected by the indicated antibodies. **C**. Crosslink between endogenous actin and AMOT was detected in cells infected with *L. pneumophila* overexpressing RavJ. HEK293T cells transfected to express the FcγII receptor were infected with the indicated bacterial strains for 4 h at an MOI of 50. Cells were lysed with 0.2% saponin and then resolved by SDS-PAGE. The blots were probed with the indicated antibodies.

To further determine the activity of this transglutaminase, recombinant RavJ purified from *E. coli* was incubated with Flag-Actin and HA-AMOT purified from HEK293T cells, and the production of a crosslink product was detected after the reactions were allowed to proceed for 2 h at 37°C (**Fig 7B**). Consistent with results from earlier experiments with cell lysates, adding His_6_-RavJ_C101A_ to the reactions did not cause crosslink between these two proteins (**Fig 7B**). Together, these results establish that RavJ is a transglutaminase that catalyzes crosslink between AMOT and actin.

### *L. pneumophila* induces crosslinking between actin and AMOT in a RavJ-dependent manner

Our results from ectopic expression by transfection strongly suggest that RavJ catalyzes protein crosslink between AMOT and actin. To determine whether this reaction is physiologically relevant, we attempted to determine the activity of RavJ during *L. pneumophila* infection. HEK293T cells transfected to express the FcγII receptor were infected with opsonized bacteria of relevant *L. pneumophila* strains at an multiplicity of infection (MOI) of 50. No crosslink between AMOT and actin was detected in cells infected with strain Lp03, an avirulent *dotA* mutant, or the wild-type stain Lp02 (**S4 Fig**). Considering the possibility that the amount of crosslinked products was too low in samples infected with strain Lp02, we examined whether overexpression of RavJ in the wild-type strain background allows us to detect the crosslink products. Indeed, infection of the cells with strain ΔravJ(pRavJ) led to robust crosslink between AMOT and actin (**Fig 7C**). In contrast, crosslink did not occur in cells similarly infected with strain ΔravJ(pRavJ_C101A_), which overexpressed the enzymatically inactive RavJ mutant (**Fig 7C**). These results indicate RavJ catalyzes crosslink between actin and AMOT in cells infected with *L. pneumophila* competent in the Dot/Icm system that overexpressed wild-type RavJ. The amount of crosslink products in cells infected with the wild-type strain likely was not sufficient for detection with our method, RavJ likely catalyzes the crosslink between actin and the Motin family of proteins.

### Simultaneous knockdown of *AMOT* and *AMOTL1* interferes with the formation of actin filaments induced by RavJ

Ectopic expression of RavJ in mammalian cells led to the formation of actin filaments (**Fig 1**). The observation that RavJ induces protein crosslink between AMOT and actin suggests that this event is important for the actin polymerization phenotype. To examine the relevance between protein crosslink and the formation of actin filaments, we determined the impact of AMOT depletion on RavJ-induced actin filament formation. To this end, we introduced shRNAs that target mRNAs of both AMOT and AMOTL1 into HEK293T cells, which is known to express high levels of these two isoforms but almost an undetectable level of AMOTL2[30]. Introduction of shRNAs by lentiviral transduction led to significant reduction of the protein levels of both AMOT and AMOTL1 (**Fig 8A**). Next, we expressed RavJ in these cells by transfection and examined the formation of actin filaments by phalloidin staining. In samples that received scramble shRNAs, actin filaments were readily detected at rates similar to untreated cells (**Fig 8B**). Consistently, crosslink between actin and AMOT became undetectable in these cells (**Fig 8C**). Furthermore, overexpression of AMOT by transfection rescued not only RavJ-induced formation of actin filaments but also the crosslink between actin and AMOT (**Fig 8B-C**). Taken together, these results support a conclusion that members of the Motin family of proteins are important for RavJ-induced actin accumulation by forming protein conjugates with actin.

**Fig 8.**
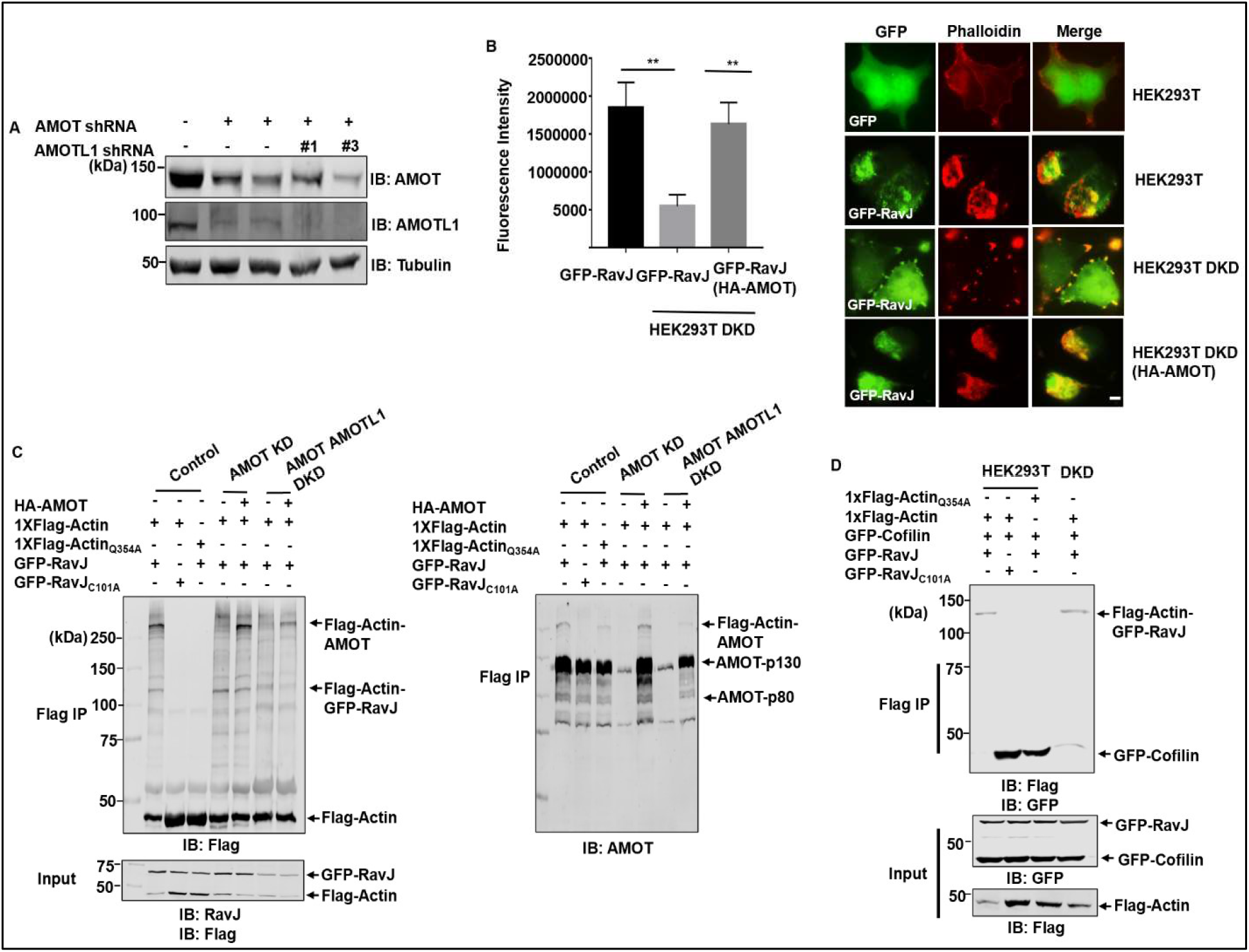
Knockdown of AMOT and AMOTL1 reduced actin accumulation in HEK293T cells induced by RavJ. **A**. Expression level of AMOT and AMOTL1 was examined by Western blot. The expression of the proteins was probed in cell lysates of two clones of AMOT knockdown cells and two clones of AMOT and AMOTL1 double knockdown cells. Proteins were detected by an AMOT-p130-specific antibody and an AMOTL1-specific antibody. Tubulin was detected as a loading control. **B**. AMOT and AMOTL1 double knockdown in HEK293T cells significantly reduced actin accumulation in cells expressing wild-type RavJ. Cells transfected to express the indicated proteins were fixed and stained with phalloidin conjugated with Texas-red. Bar, 10 μm. Phalloidin fluorescence signal was quantified by Image J software to evaluate the amount of F-actin (the left panel). Data are the mean SEM (***p<0.001). **C**. Knockdown of AMOT and AMOTL1 in HEK293T cells abolished crosslink between actin and the Motin family of protein. Cells were transfected to express the indicated proteins and samples were subjected to IP with beads coated with the Flag antibody. Crosslinking products were detected with the Flag-specific antibody and AMOT-specific antibodies, respectively. **D**. RavJ inhibits the binding between actin and cofilin. Cells transfected to express the indicated proteins were lysed and subjected to IP with beads coated with the Flag-specific antibody. Binding between actin and cofilin were detected by the indicated antibodies.

### RavJ interferes with the interaction between actin and cofilin

Earlier studies suggest that actin interacts with profilin and cofilin through its Gln_354_ residue[31–33]. Cofilin depolymerizes and severs actin filaments while profilin binds to actin monomers and provides ATP-Actin for incorporation into actin filaments[34]. The fact that actin crosslinks with RavJ and AMOT via Gln_354_ inspired us to investigate whether RavJ affects the binding between actin and actin binding proteins. HA-profilin-1, HA-profilin-2 or GFP-cofilin were expressed in HEK293T cells along with relevant proteins, and cell lysates were subjected to immunoprecipitation with the Flag-specific antibody. Interestingly, expression of RavJ reduced the binding between actin and cofilin (**Figure 8D**), but not profilin-1 and profilin-2 (**S5A-B Fig**), suggesting that RavJ-induced actin accumulation in cell cortex was a result of reduced actin depolymerization.

### LegL1 blocks the activity of RavJ by sterically hindering its catalytic site

In a previous study, LegL1 (Lpg0945) was identified as a putative metaeffector of RavJ which interacts with its amino terminus containing the predicted motif important for its enzymatic activity[27]. We then tested whether LegL1 can influence actin cytoskeleton rearrangement induced by RavJ. Coexpression of HA-LegL1 with GFP-RavJ significantly reduced the formation of crosslink products by actin and AMOTs and such reduction became more apparent as the ratio between LegL1 and RavJ increased (**Fig 9A**), indicating that LegL1 inhibits the activity of RavJ. The inhibitory effect of LegL1 can attribute to at least two potential mechanisms: LegL1 reverses the crosslink by an enzymatic activity, or it inhibits the activity of RavJ by direct binding.

**Fig 9.**
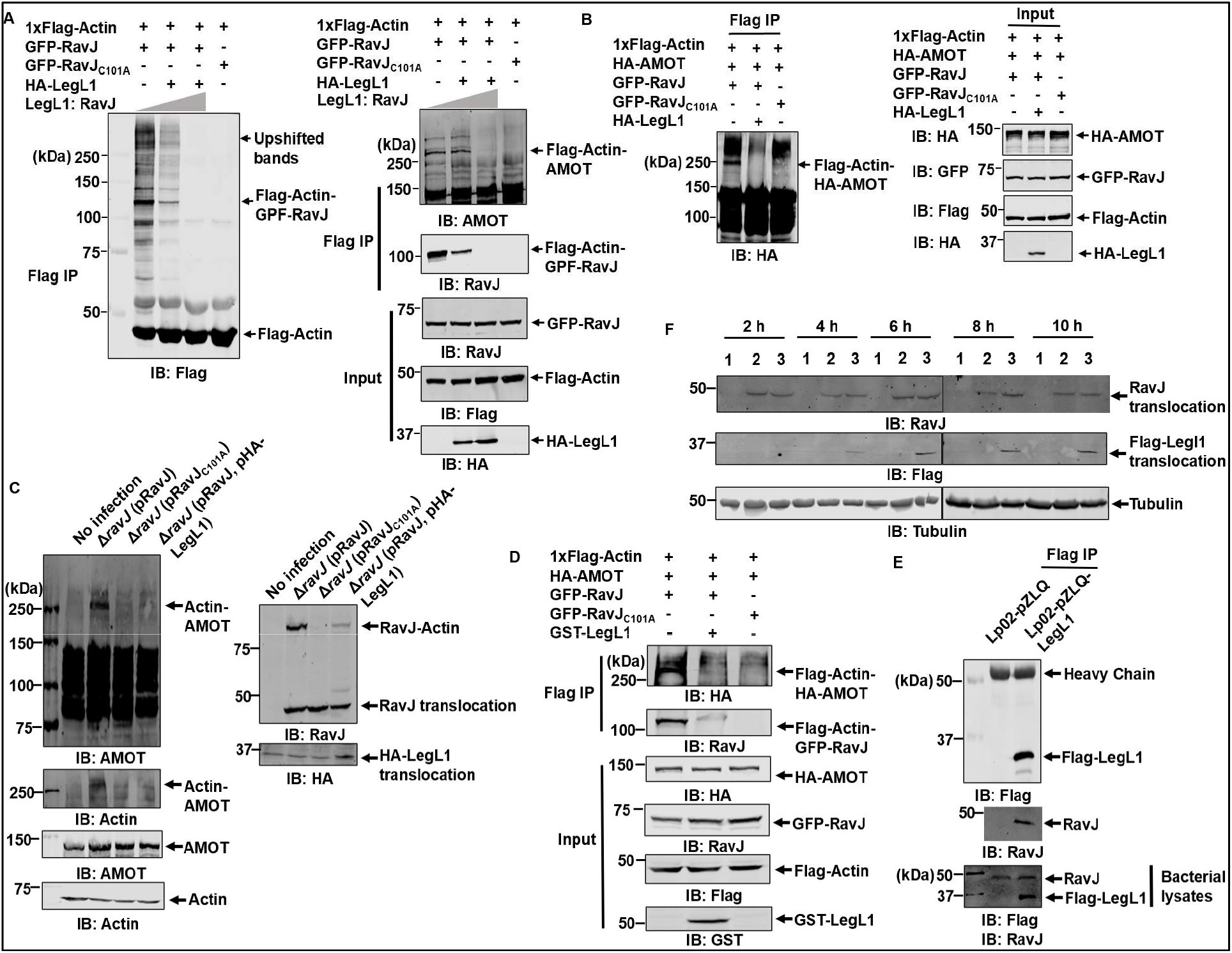
LegL1 antagonizes the catalytic activity of RavJ. **A**. expression of LegL1 in HEK293T cells reduced crosslink products induced by RavJ. Lysates of cells transfected to express the indicated proteins were subjected to IP with beads coated with the Flag antibody and the products resolved by SDS-PAGE were detected with the indicated antibodies. **B**. LegL1 inhibits the activity of RavJ in a cell-free system. Cells transfected to express Flag-Actin, HA-LegL1, HA-AMOT, GFP-RavJ, and GFP-RavJ_C101A_, respectively, were lysed with RIPA buffer without EDTA. Lysates of HA-LegL1 was pre-incubated with those expressing GFP-RavJ for 1 h at 37°C. Reactions containing the indicated cell lysates were immunoprecipitated with beads coated with the Flag antibody and proteins were detected with the indicated antibodies. **C**. Overexpression of LegL1 in *L. pneumophila* inhibits the enzymatic activity of RavJ. Bacteria of the indicated *L. pneumophila* strains were opsonized prior to infecting HEK293T cells transfected to express the FcγII receptor at an MOI of 50 for 4 h. Cells lysed with 0.2% saponin were probed with the indicated antibodies. **D**. Recombinant LegL1 reduces the crosslink products induced by RavJ. Cells transfected to express Flag-Actin, HA-AMOT, GFP-RavJ, and GFP-RavJ_C101A_, respectively, were lysed with RIPA buffer without EDTA. 10 μg GST-LegL1 was pre-incubated with cell lysates of GFP-RavJ for 1 h at 37°C. Reactions containing the indicated components were immunoprecipitated with beads coated with the Flag antibody and proteins were detected with the indicated antibodies. **E**. LegL1 directly binds to RavJ in *L. pneumophila*. Bacterial cells carrying either the vector pZLQ or pZLQ-LegL1 were lysed with RIPA buffer and then immunoprecipitated with beads coated with the Flag antibody. Binding between RavJ and LegL1 was detected by RavJ-specific antibodies and the Flag antibody, respectively **F**. Overexpression of LegL1 in *L. pneumophila* does not influence the translocation of RavJ. HEK293T cells transfected to express the FcγII receptor were treated with different bacterial strains: 1. No infection, 2. Wildtype *L. pneumophila*. 3. Wild type *L. pneumophila* overexpressing Flag-LegL1. Cells were collected at the indicated time points and lysed by 0.2% saponin. RavJ and LegL1 translocation were detected by RavJ-specific antibodies and the Flag-specific antibody respectively. Tubulin was probed as a loading control.

To distinguish between these two possibilities, we pre-incubated cell lysates expressing HA-LegL1 with cell lysates expressing GFP-RavJ for 1 h at 37°C. Cell lysates expressing HA-AMOT and Flag-Actin, respectively, were then added to the cell-free system. RavJ that has been pre-incubated with HA-LegL1 was unable to induce crosslink between actin and AMOT, suggesting that LegL1 blocks the activity of RavJ (**Fig 9B**). Furthermore, addition of HA-LegL1 to reactions containing crosslinked actin-AMOT did not detectably reduce the amount of the conjugate product (**S6 Fig**). We also tested the activity of recombinant LegL1 purified from *E. coli*. Incubation of GST-LegL1 with cell lysates expressing GFP-RavJ ablated its ability to catalyze crosslink between actin and AMOT (**Fig 9D**).

We next examined the interactions between LegL1 and RavJ in *L. pneumophila*. RavJ can be immunoprecipitated from lysates of *L. pneumophila* expressing Flag-Leg1 with beads coated with the Flag antibody, indicating binding between LegL1 and RavJ (**Fig 9E**). We further tested the effect of LegL1 on RavJ-induced crosslink during bacterial infection. HEK293T cells were infected with *ravJ* mutant complemented with RavJ, the RavJ_C101A_ mutant, or both RavJ and LegL1, at an MOI of 50. Crosslink between AMOT and actin was only detected in cells infected with the ΔravJ(pRavJ) strain but not strain ΔravJ(pRavJ, pLegL1) (**Fig 9C**), suggesting that LegL1 effectively inhibits the crosslink caused by RavJ during bacterial infection. The translocation of RavJ into host cells is not influenced by LegL1, indicating that the regulation of RavJ by LegL1 is likely occurs in the host cells (**Fig 9F**). Collectively, these results indicate that LegL1 blocks the activity of RavJ by direct binding.

## Discussion

The actin cytoskeleton network is a major host structural component that provides structural and functional support in numerous vital cellular activities. It also directs the trafficking of cargo-containing vesicle trafficking throughout the cell by functioning as a highway[35]. Intracellular bacterial pathogens have evolved remarkable strategies to subvert the host cytoskeletal machinery to promote bacterial internalization, facilitate the biogenesis of bacteria-containing phagosomes, and co-opt actin-dependent movement to benefit pathogen dissemination[36]. To support diverse infection events, intracellular pathogens hijack the actin cytoskeleton by introducing effector proteins into the host cytosol by specialized secretion systems[8]. Among these, *Coxiella burnetii* triggers actin reorganization at the attachment site of phagocytic human macrophages by binding to CR3 receptors to stimulate bacterial internalization[37]. After entering a host cell, this bacterium replicates within an acidic compartment called the parasitophorous vacuole (PV) in macrophages. Optimal intracellular growth of *C. burnetii* requires F-actin accumulation around the PV, but the detailed mechanisms are not fully characterized[38]. *Chlamydia* species replicate in a host membrane-derived compartment termed inclusion. Optimal development and maintenance of vacuole morphology and integrity require F-actin rings surrounding the inclusion to stabilize the organelle[39, 40]. Spotted Fever Group *Rickettsia*, such as *R. rickettsii* and *R. conorii*, escape the phagosome before lysosomal fusion after entering a host cell. After escape, these bacteria induce actin polymerization to form an actin tail that facilitates bacterial motility within the cell[41, 42].

Actin exists in two main forms of organization, the monomeric G-actin, and the filamentous form F-actin[43]. Polymerization and depolymerization of actin filaments are kept in a dynamic balance to tightly regulate movement and other cell functions[43]. The VCA domains of N-WASP (Wiskott-Aldrich syndrome protein) and SCAR/WAVE (suppressor of Camp receptor/WASP-family verprolin homologous protein) activate the actin nucleator Arp2/3 complex to generate new F-actin branches from preexisting mother filaments[44]. Actin-depolymerizing factor ADF/cofilin regulates actin dynamics by depolymerizing filaments at their pointed ends, thereby restoring a pool of actin monomers for filament assembly[35]. Actin molecule harbors two main lobes separated by a deep upper cleft. Each main lobe is subdivided into two clearly discernible subdomains, SD1-4[45]. SD1, which is built from residues 1-32, 70-144, and 338-375, forms an important target area for a large number of actin-binding proteins such as profilin, cofilin and gelsolin[45]. Of note is that RavJ-induced actin crosslink with AMOT occurs at Gln_354_, a site locates on the SD1, suggesting that this crosslinking event may interfere with the interaction between actin subunits in the filament and the actin binding proteins. In agreement with this notion, the binding between cofilin and actin is significantly reduced (**Fig 8D**), indicating that RavJ functions to block actin depolymerization, resulting in the accumulation of F-actin in cell cortex.

It has been known for a long time that *L. pneumophila* avoids the delivery of its vacuole to lysosomes by modulating the ER-to-Golgi vesicle trafficking[46]. Shortly after being internalized by a host cell, the plasma membrane derived vacuole containing *L. pneumophila* will be converted into a compartment that has similarity to an ER-Golgi intermediate compartment[47–49]. The biogenesis of this specialized phagosome has been studied extensively[49]. A repertoire of Dot/Icm effectors have been demonstrated to hijack the host vesicle trafficking pathway directly[2, 50], some of them could affect this cellular process indirectly to bypass the microbicidal endosomal compartment[51]. The modulation of the actin cytoskeleton clearly contributes to the development of the LCV. For example, the metal protease RavK cleaves actin, abolishing its ability to form actin polymers[11] whereas LegK2 phosphorylates components of actin nucleator ARP2/3 complex and thus inhibits actin polymerization on the phagosome[11]. Ceg14 inhibits actin polymerization by a yet unknown mechanism[13]. These proteins are very likely working in synergic to temporally inhibit actin polymerization on the LCV and thus preventing fusion with late endosomes[12]. In contrast to the inhibitory effects of RavK and LegK2, VipA enhances actin polymerization by acting as an actin nucleator[14]. RavJ also appears to promote the formation of actin filaments. Here, we showed that RavJ-induced crosslink between actin and AMOT blocks the depolymerization activity of ADF/cofilin, resulting in the stabilization of actin polymers in cell cortex (**S7 Fig**). Clearly, in cells infected with *L. pneumophila*, its effectors strike a balance between the two states of actin. Balanced modulation of host processes has recently emerged as a prominent feature associated with the interactions between *L. pneumophila* and its hosts. In some cases, the balance is achieved by effector pairs with opposite biochemical activity which may function at different phases of infection[52]. In other cases, the importance of balance lies in the cellular locations of the molecular events regulated by the effectors[17]. In addition to other effectors that inhibit actin polymerization, the regulation imposed by RavJ is controlled by its metaeffector LegL1, which blocks its function by direct protein-protein interactions. LegL1 likely functions to spatially and temporally prevent RavJ from inducing excessive actin polymerization.

Clathrin-vesicle associated proteins regulate vesicle assembly by binding directly to the actin filaments through a C-terminal talin-like domain, indicating important correlation between actin polymerization and endocytic vesicle trafficking[53]. Actin filaments also play a key role in maintaining ER structure and the ER-to-Golgi trafficking[54]. Given the extensive interconnection between vesicle trafficking and the actin cytoskeleton, RavJ is likely working in concert with VipA to enhance cargo transportation between ER and the LCV, thus facilitating fusions between the ER-derived vesicles and the LCV. Together, these effectors are likely work in concert to promote bacterial replication by indirectly interfering with the vesicle trafficking pathway.

## Materials and Methods

### Media, bacteria strains, and cell lines

*E. coli* strain DH5a was used for cloning and plasmid construction strains XL1-Blue and BL21(DE3) were used for expression and purification of all the recombinant proteins used in this study. *E. coli* strains were grown on LB agar plates or in LB broth. When necessary, antibiotics were added to media at the following concentrations: ampicillin, 100 μg/ml; kanamycin, 30 μg/ml. *L. pneumophila* strains used in this study were derivatives of the Philadelphia 1 strain Lp02[55]. Lp03 is an isogenic *dotA^-^* mutant of Lp02[56]. All strains were grown and maintained on CYE plates or in ACES-buffered yeast extract (AYE) broth as previously described[55]. For *L. pneumophila*, kanamycin was used at 30 μg/ml. When needed, thymidine was added at a final concentration of 100 μg/ml for thy autotrophic strains. The *ravJ* in-frame deletion strain was constructed by a two-step allelic exchange strategy as described previously[57]. HEK293T cells purchased from ATCC were cultured in Dulbecco’s modified minimal Eagle’s medium (DMEM) supplemented with 10% fetal bovine serum (FBS). Bone marrow-derived macrophages (BMDMs) were isolated and cultured as described previously[58]. All cell lines were regularly checked for potential mycoplasma contamination by the universal mycoplasma detection kit from ATCC (cat# 30-1012K).

### Plasmid constructions

All the plasmids used in this study are listed in Table S1 and the bacterial strains and antibodies are in Table S2. For protein purifications, *ravJ, ravJ_C101A_*, and *legl1* were cloned into pQE30 (QIAGEN) or pGEX6p-1, respectively. For complementation experiments, *ravJ* and *ravJ_C101A_* were inserted into pZLQ-Flag, a derivative of pZLQ[59] that was modified to carry a Flag tag[60]. *legl1* was inserted into either pZLQ-Flag or pZL507[61] for overexpression in *L. pneumophila*. For ectopic expression of proteins in mammalian cells, genes were inserted into the 4xFlag CMV[20], pFlag-CMV (Sigma), pEGFP-C1 (Clontech) vector, 3XHA pCDNA3.1 vector[62] or pAPH[63], a derivative of pVR1012 suitable for expressing proteins with an amino terminal HA tag. Human *AMOT* and *AMOTL1* were amplified from cDNAs of HEK293T cells and then inserted into BamHI/SalI of pAPH. For shRNA knockdown of *AMOTL1* in mammalian cells, pLKO. 1 -hygro vector (Addgene, plasmid #24150) was used to generate the pLKO.1-hygro-AMOTL1-sh construct. Packing plasmids psPAX2(Addgene, plasmid #12260) and pMD2.G (Addgene, plasmid #12259) were used for lentiviral constructs transduction.

### Transfection, immunoprecipitation, infection

HEK293T cells grown to about 90% confluence were transfected with different plasmids, respectively, using Lipofectamine 3000 (Thermo Fisher Scientific). Transfected cells were collected and lysed with radioimmunoprecipitation assay buffer (RIPA buffer, Thermo Fisher Scientific) at 18-24 h post transfection. When needed, immunoprecipitation was performed with lysates of transfected cells with Flag- or HA-specific antibody-coated agarose beads (Sigma-Aldrich, cat# F2426, Pierce, cat# 88836, respectively) at 4°C overnight. Beads were then washed with pre-cold RIPA buffer three times. For tandem purification, followed by RIPA wash, agarose beads with flag-tagged proteins were washed with Flag-to-His buffer (100 mM Na-Phosphate, pH 8.0, 150 mM NaCl, 0.05% Triton X-100) three times and eluted with 3XFLAG peptide (Sigma-Aldrich, cat# F3290). Elution fraction was then subjected to HA beads, or Ni^2+^-NTA agarose beads (QIAGEN) as needed. Beads were resolved by SDS-PAGE gels followed by immunoblotting analysis with specific antibodies or silver staining following the manufacturer’s protocols (Sigma-Aldrich, cat# PROTSIL1).

For infection experiments, *L. pneumophila* strains were grown to post-exponential phase (OD_600_=3.2-3.8) in AYE broth. Complementation strains and overexpression strains were induced with 0.5 mM IPTG for 2 h at 37°C before infection. HEK293T cells were transfected to express FcγII receptor[20]. *L. pneumophila* strains were incubated with *L. pneumophila-specific* anti-sera at a dilution of 1:500 for 30 minutes at 37°C. Cells were infected at an MOI of 50 for 4 h and then lysed with 0.2% saponin. Cell lysates were resolved by SDS-PAGE and followed by immunoblotting analysis with the specific antibodies.

### Antibodies and immunoblotting

Purified His_6_-RavJ was used to raise rabbit specific antibodies following a standard protocol (Pocono Rabbit Farm & Laboratory). The antibodies were affinity-purified as described before[64]. For immunoblotting, samples resolved by SDS-PAGE were transferred onto 0.2 μm nitrocellulose membranes (Bio-Rad, cat# 1620112). Membranes were blocked with 5% non-fat milk, incubated with the appropriate primary antibodies: anti-HA (Sigma-Aldrich, cat# H3663), 1:5000; anti-tubulin (DSHB, E7), 1:10000, anti-Flag (Sigma-Aldrich, cat# F1804), 1:5000, anti-RavJ (this study), 1:5000, anti-Actin (MP Biomedicals, cat# 0869100), 1:5000, anti-AMOTL1 (Sigma-Aldrich, cat# SAB1408393), 1:5000, anti-AMOT (Abnova, cat# H00154796-B01P), 1:5000, anti-ICDH[61], 1:10000, anti-GFP[61], 1:10000, anti-GST[61]. Membranes were then incubated with an appropriate IRDye infrared secondary antibody and scanned by an Odyssey infrared imaging system (Li-Cor’s Biosciences).

### Immunostaining

HEK293T cells were seeded at 1X10^5^ per well on glass coverslips in 24-well plates. Cells were transfected to express the corresponding proteins for 24 h and were washed three times with PBS. Cells were fixed with 4% formaldehyde for 30 minutes at room temperature, washed with PBS three times and were then permeabilized by 0.3% Triton at room temperature for 15 minutes. F-actin was stained with phalloidin conjugated with Texas-red (Thermo Fisher Scientific, cat#T7471) at a dilution of 1:500 for 1 h at room temperature. Images were acquired using an Olympus X-81 fluorescence microscope.

### Protein purification

10 ml overnight *E. coli* cultures were transferred to 400 ml LB medium supplemented with 100 μg/ml ampicillin or 30 μg/ml kanamycin and grown to OD_600nm_ of 0.8-1.0. Cultures were then incubated at 18°C for 16-18 h after the addition of IPTG at a final concentration of 0.5 mM. Bacterial cells were spun down at 12,000 g and lysed by sonication. The soluble lysates were cleared by spinning at 12,000g twice at 4°C for 20 minutes. To purify His_6_-tagged proteins, supernatants were incubated with Ni^2+^-NTA beads for 2 h at 4°C followed by elution with 300 mM imidazole in TBS buffer after washing with 20X bed volumes of TBS buffer containing 20 mM imidazole. Purified proteins were dialyzed in buffer containing TBS, 5% glycerol and 1 mM DTT overnight at 4°C. GST-tagged proteins were purified with glutathione beads (Pierce, cat# 16101) for 2 h at 4°C followed by elution with elution buffer (50 Mm Tris pH 8.0, 0.4 M NaCl, 50 mM reduced glutathione, 0.1% Triton X-100, 1 mM DTT).

### shRNA knockdown of *AMOT* and *AMOTL1*

MISSION shRNA retroviral constructs targeting *AMOT* was purchased from Sigma-Aldrich (Clone ID: NM_133265.1-1628s1c1). To collect viral supernatant, cells were seeded at 2×10^5^ cells/6 cm plate (around 10% confluency). After overnight culture, cells were transfected with the retroviral construct targeting *AMOT* along with two packing plasmids, psPAX2 and pMD2.G. The viral supernatant was collected at day 5 after transfection and was filtered with a 0.45-μm syringe filter. The titer of the produced lentivirus was determined by using Lenti-X Gostix Plus Titer Kit (Takara, cat# 631281). To generate *AMOT* knockdown cell line, HEK293T cells were seeded at 2×10^5^ cells/10 cm plate (around 10% confluency). Media was removed at day 2 and viral supernatant was added to cover the whole plate (3.5 ml/10 cm plate). Viral supernatant was added every 3 hours for three times. At day 3, media was replaced with fresh media. Cells were selected with media supplemented with puromycin (InvivoGen, cat# ant-pr-1) for a few days and single colonies were selected.

To generate *AMOTL1* knockdown cell line, the pLKO.1-hygro-*AMOTL1* -sh construct was generated by cloning annealed oligos 5’-CCGGTCCGGGCCCATCCTACAAACAACTTTCTCGAGAAAGTT GTTTGTAGGATGGGCTTTTTTGG-3’ and 5’-AATTCCAAAAAAGCCCATCCTACAAACAACTTTCTCGAGAAAGTTGTTTGTAGGATGGGCCCGGA-3’ into the pLKO.1-hygro vector. Viral supernatant was generated by transfecting HEK293T cells with the retroviral construct targeting *AMOTL1* along with the packing plasmids, psPAX2 and pMD2.G. The viral supernatant was collected as described above. The *AMOT* knockdown cell line was infected with the viral supernatant as described above and single colonies were selected with media supplemented with hygromycin (Thermo Fisher Scientific, cat#10687010).

### Cell-free assays and *in vitro* assays

In the cell-free assays, HEK293T cells expressing Flag-Actin, HA-AMOT, GFP-RavJ, GPF-RavJ_C101A_, HA-LegL1, respectively, were lysed by RIPA buffer without EDTA. Cell lysates were spun down at 12,000 g and the supernatants were collected. Reactions with combined supernatants as indicated were allowed to proceed for 2h at 37°C. the supernatants were then subjected to immunoprecipitation with Flag-antibody coated beads or HA-antibody coated beads as needed.

For *in vitro* assays, HEK293T cells expressing HA-AMOT, Flag-Actin respectively were lysed with RIPA buffer and then subjected to HA-IP or Flag-IP as needed. Proteins were eluted from the corresponding beads using 3XFLAG peptide or HA peptide (Thermo Fisher Scientific). 5 μg His_6_-RavJ or His_6_-RavJ_C101A_ were added to the *in vitro* assay and the reaction was left in the 37°C incubator for 2 h. The in vitro reaction was then subjected to Flag-IP followed by SDS-PAGE analysis.

### Intracellular bacterial growth assay

*L. pneumophila* strains were grown to the post-exponential phase (OD_600_=3.2-3.8) before infection. Bone marrow-derived mouse macrophages (BMDMs) isolated from female A/J mice as described before[58] were seeded onto 24-well plates and were infected with relevant *L. pneumophila* strains at an MOI of 0.05 at 37°C. Cells were collected at the indicated time points and lysed with 0.02% saponin for half an hour on ice. The bacteria number was determined by enumerating colony-forming unit (CFU) of appropriately diluted saponin-soluble fractions.

### LC-MS/MS analysis

Protein bands were digested in-gel with trypsin for protein identification. Peptides were re-suspended in 96.9% water, 3% acetonitrile (ACN), and 0.1% formic acid (FA) at the final concentration of 0.2μg/μl, and 1 μg total peptides (equivalent volume) was analyzed by LC-ESI-MS/MS system using the Dionex UltiMate 3000 RSLC nano System coupled to the Q Exactive™ HF Hybrid Quadrupole-Orbitrap Mass Spectrometer (Thermo Scientific, Waltham, MA) as described previously[65, 66]. The reverse phase peptide separation was accomplished using a trap column (300 μm ID × 5mm) packed with 5 μm 100 Å PepMap C18 medium, and then separated on a reverse phase column (50-cm long × 75μm ID) packed with 2 μm 100 Å PepMap C18 silica (Thermo Fisher Scientific, Waltham, MA). The column temperature was maintained at 50 °C.

Mobile phase solvent A was 0.1% FA in water and solvent B was 0.1% FA in 80% ACN. Loading buffer was 98%/water/2% ACN/0.1% FA. Peptides were separated by loading into the trap column in a loading buffer for 5-min at 5 μL/min flow rate and eluted from the analytical column at a flow rate of 150 nL/min using a 130-min LC gradient as follows: linear gradient of 5.1 to 27% of solvent B in 80 min, 27-45% in next 20 min, 45-100% of B in next 5 min at which point the gradient was held at 100% of B for 7 min before reverting back to 2% of B at 112 min, and hold at 2% of B for next 18 min for equilibration. The mass spectrometer was operated in positive ion and standard data-dependent acquisition mode with Advanced Peak Detection function activated for the top 20n. The fragmentation of precursor ion was accomplished by stepped normalized collision energy setting of 27%. The resolution of Orbitrap mass analyzer was set to 120,000 and 15,000 for MS1 and MS2, respectively. The full scan MS1 spectra were collected in the mass range of 350-1,600 m/z, with an isolation window of 1.2m/z and a fixed first mass of 100 m/z for MS2. The spray voltage was set at 2 and Automatic Gain Control (AGC) target of 4e5 for MS1 and 5e4 for MS2, respectively.

For protein identification, the raw data were processed with the software MaxQuant[67] (version 1.6.3.3) against *Homo sapiens* database downloaded from the UniProt (www.uniprot.org). The following parameters were edited for the searches: precursor mass tolerance of 10 ppm; enzyme specificity of trypsin enzyme allowing up to 2 missed cleavages; oxidation of methionine (M) as a variable modification and carbamidomethylation of cysteine (C) as a fixed modification. False discovery rate (FDR) of peptide spectral match (PSM) and protein identification was set to 0.01. The unique plus razor peptides (non-redundant, non-unique peptides assigned to the protein group with most other peptides) were used for peptide quantitation. LFQ intensity values were used for relative protein abundance measurement. Only proteins detected with at least one unique peptide and MS/MS ≥2 (spectral counts) were considered as true identification and used for downstream analysis.

---

**S1 Fig.**
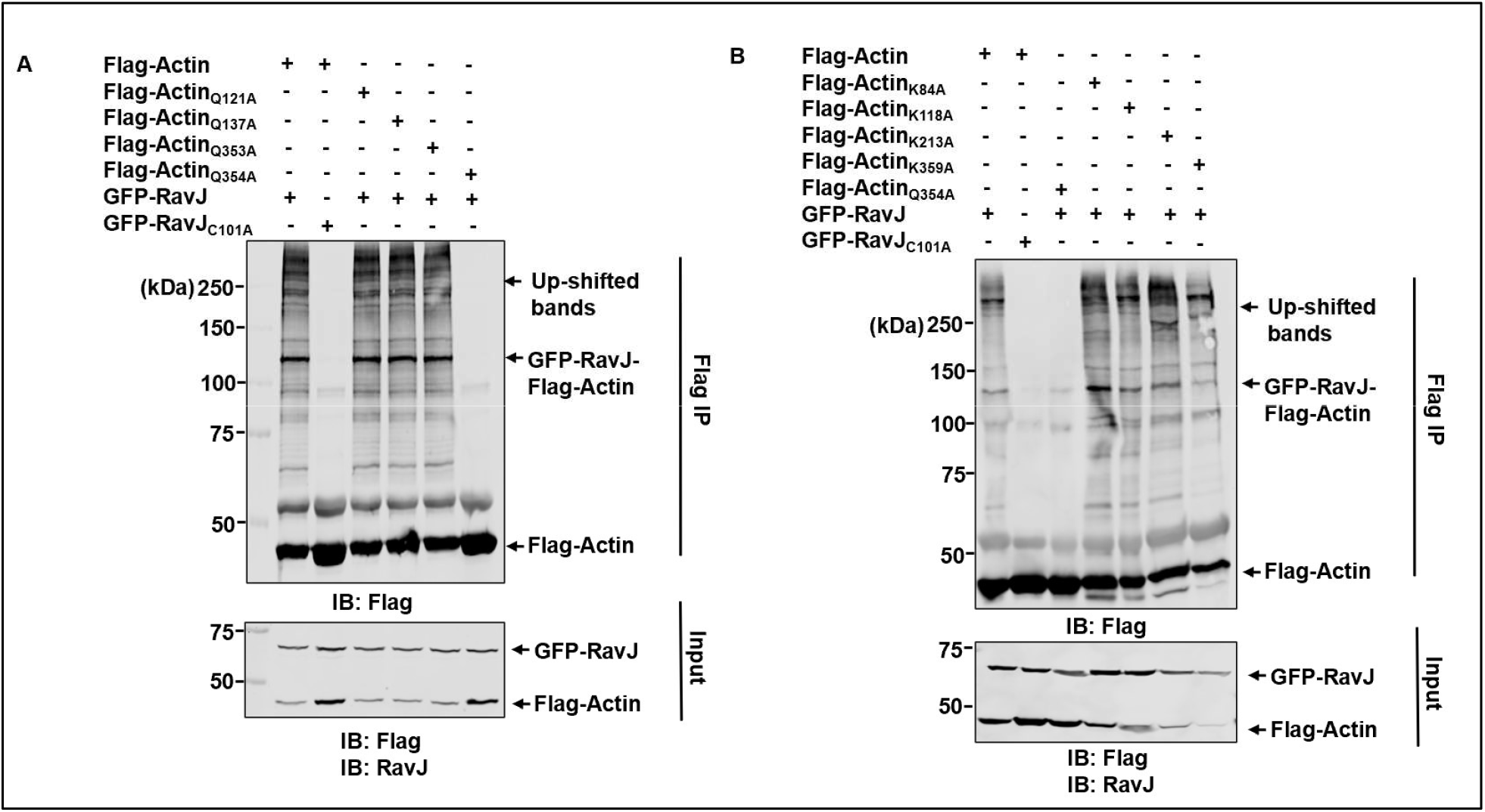
Determination of the crosslink site in actin. **A-B.** HEK293T cells co-transfected to express the indicated proteins were lysed and immunoprecipitated by beads coated with the Flag-specific antibody. Note that only the Gln_354_Ala mutation in actin abolished the crosslink products induced by RavJ.

**S2 Fig.**
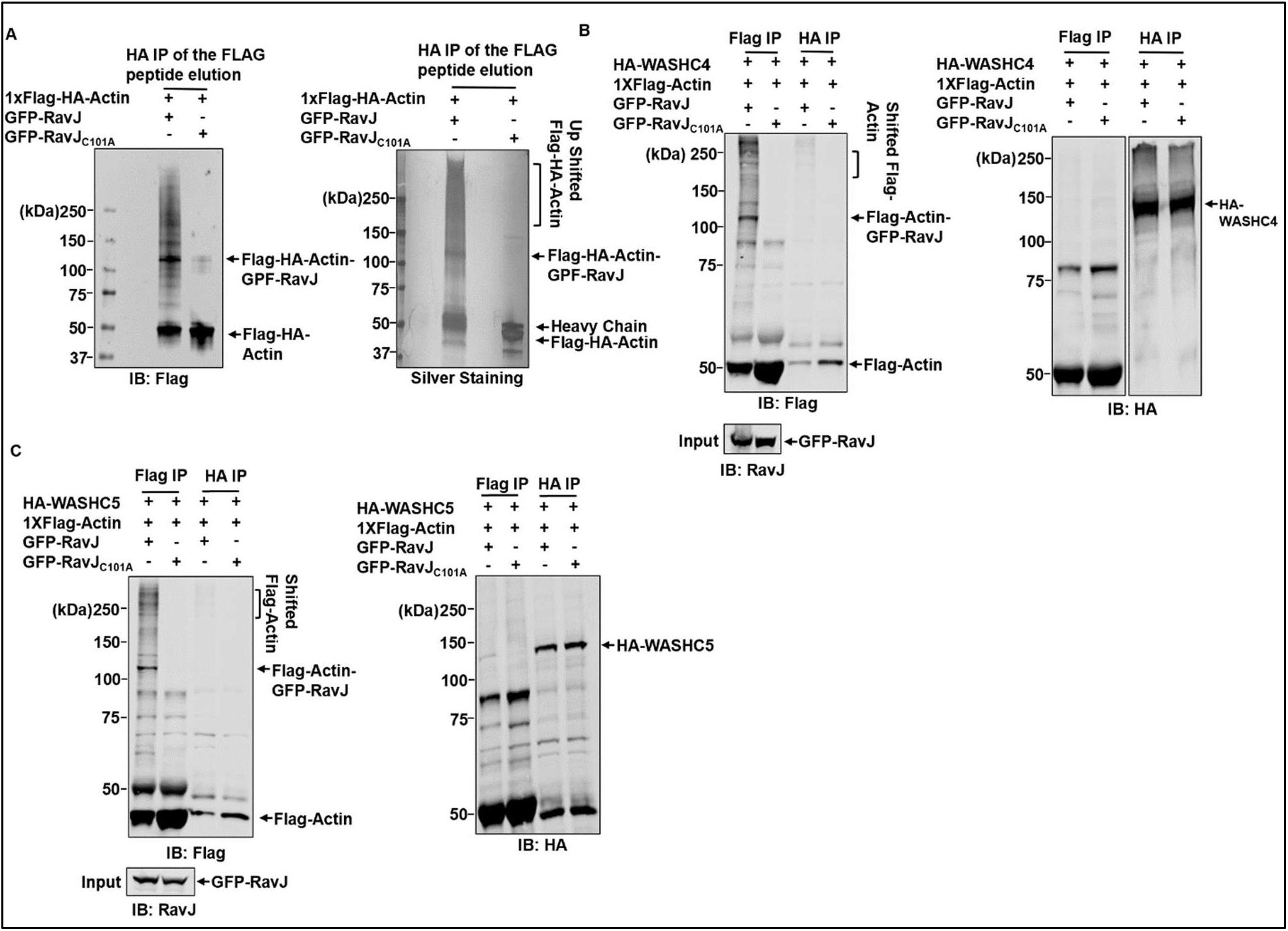
Identification of the cellular targets of RavJ. **A**. Tandem purification of actin crosslink products catalyzed by RavJ. HEK293T cells co-transfected to express Flag-HA-Actin, GFP-RavJ or GFP-RavJ_C101A_ were subjected to IP with beads coated with the Flag antibody. Proteins were eluted from the beads by 3XFLAG peptide. Elution fractions were then immunoprecipitated with beads coated with the HA-specific antibody. Products resolved by SDS-PAGE were detected by silver staining (right) or probed with the Flag antibody(left). Gel plug corresponding to the upshifted band was cut for MS analysis. **B-C**. WASHC4 and WASHC5 do not crosslink with actin in the presence of RavJ. HEK293T cells co-transfected to express the indicated proteins were lysed and immunoprecipitated by beads coated with the Flag-specific or HA-specific antibody. Note that there is no crosslink products detected between actin and WASHC4 or WASHC5.

**S3 Fig.**
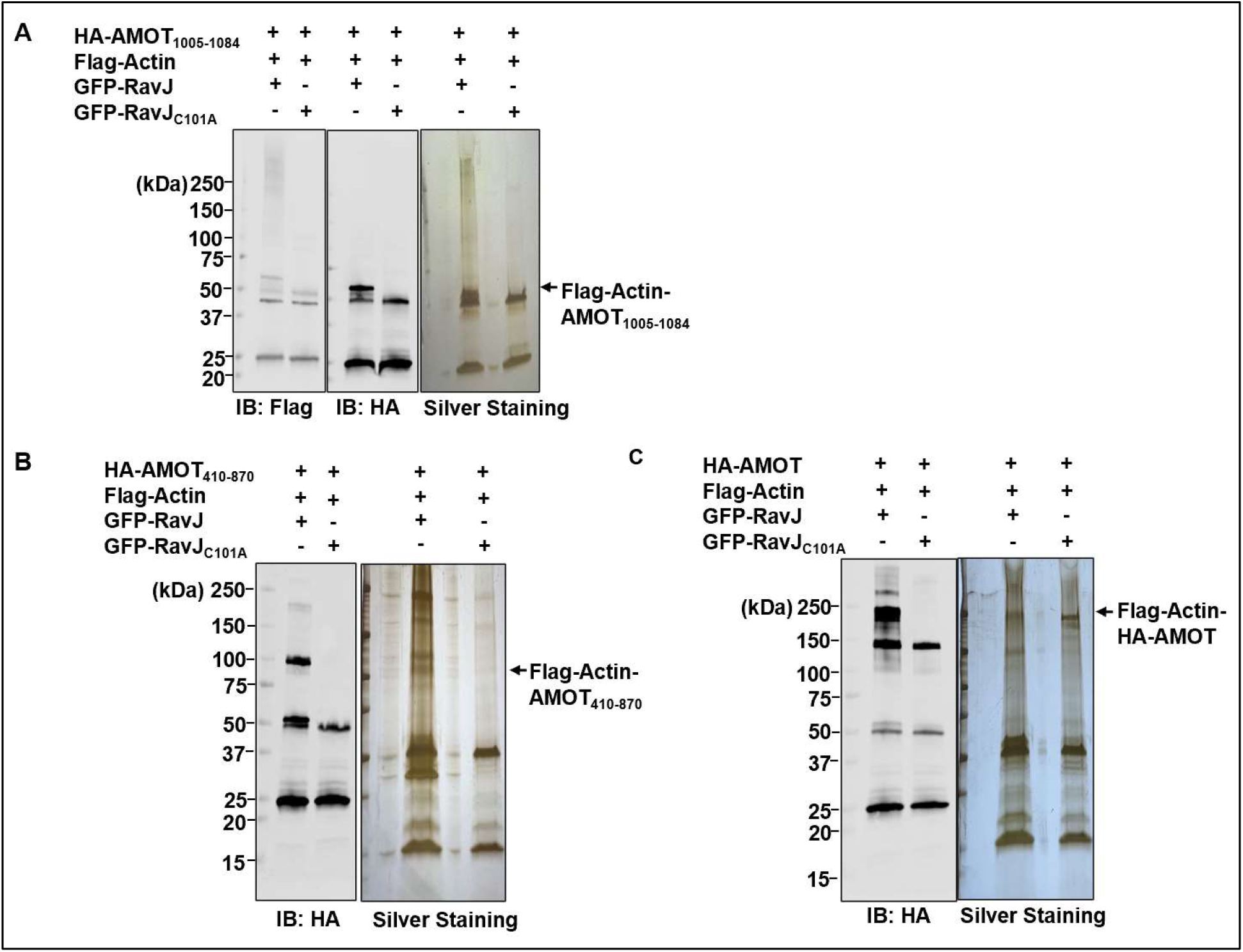
Determination of the crosslink sites in AMOT. **A-C**. Tandem purification of crosslink products of actin and AMOT truncations. HEK293T cells co-transfected to express the indicated proteins were subjected to IP with beads coated with the Flag antibody. Proteins were eluted from the beads by 3XFLAG peptide. Elution fractions were then immunoprecipitated with beads coated with the HA-specific antibody. Products resolved by SDS-PAGE were detected by silver staining (right) or probed with the indicated antibodies(left). Protein bands corresponding to the crosslink products were excised for MS analysis.

**S4 Fig.**
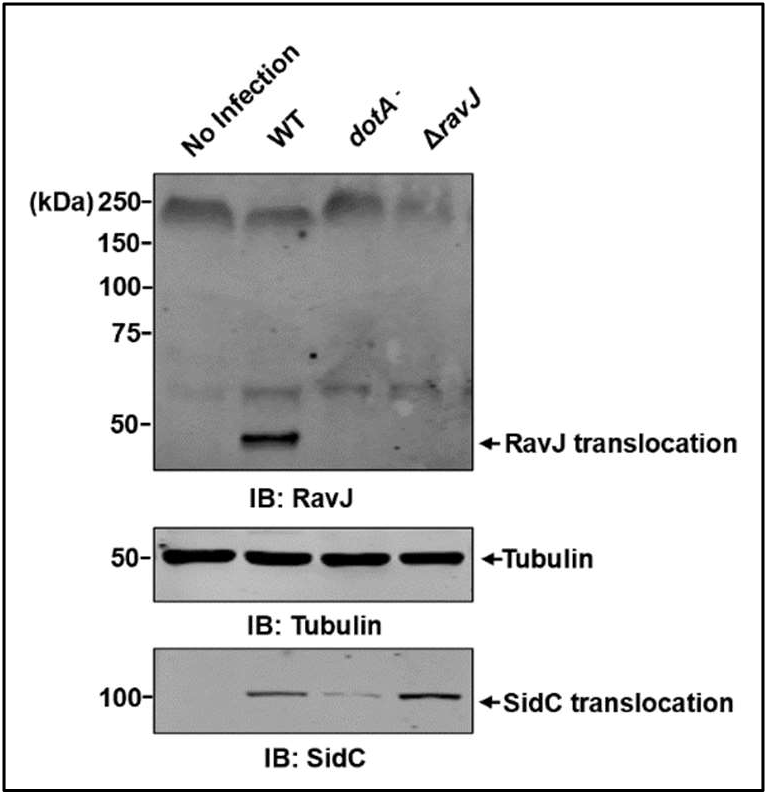
Determination of the effect of RavJ during bacterial infection. HEK293T cells transfected to express the FcγII receptor were treated with opsonized bacterial strains as indicated. Cells were collected at 4 h after infection and were lysed by 0.2% saponin. RavJ and SidC translocation were detected by RavJ-specific or SidC-specific antibodies. Tubulin was probed as a loading control.

**S5 Fig.**
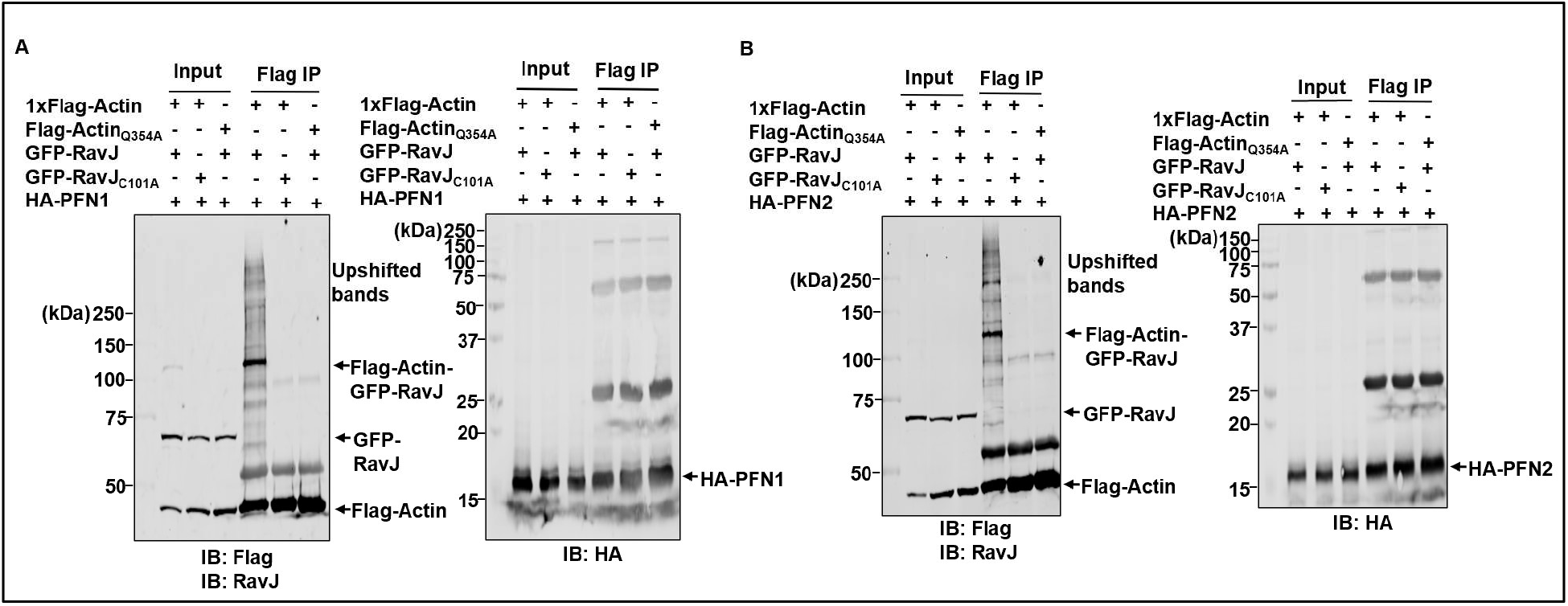
Effects of RavJ on the binding between actin and actin binding proteins. **A-B** RavJ does not influence the binding between actin and actin binding proteins, profilin 1 and profilin 2. HEK293T cells co-transfected to express the indicated proteins were lysed and subjected to IP with beads coated with a Flag-specific antibody.

**S6 Fig.**
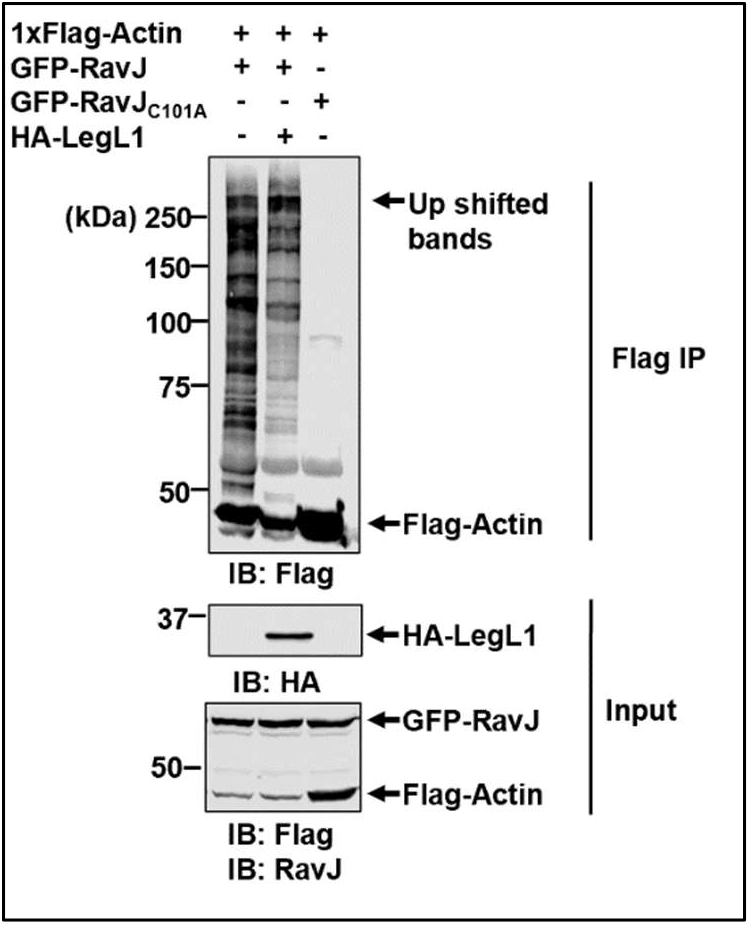
LegL1 does not reverse the transglutaminase activity of RavJ. HEK293T cells transfected to express the indicated proteins were lysed and subjected to IP with beads coated with the Flag-specific antibody. Note that expression of HA-LegL1 did not remove actin from the crosslink product.

**S7 Fig.**
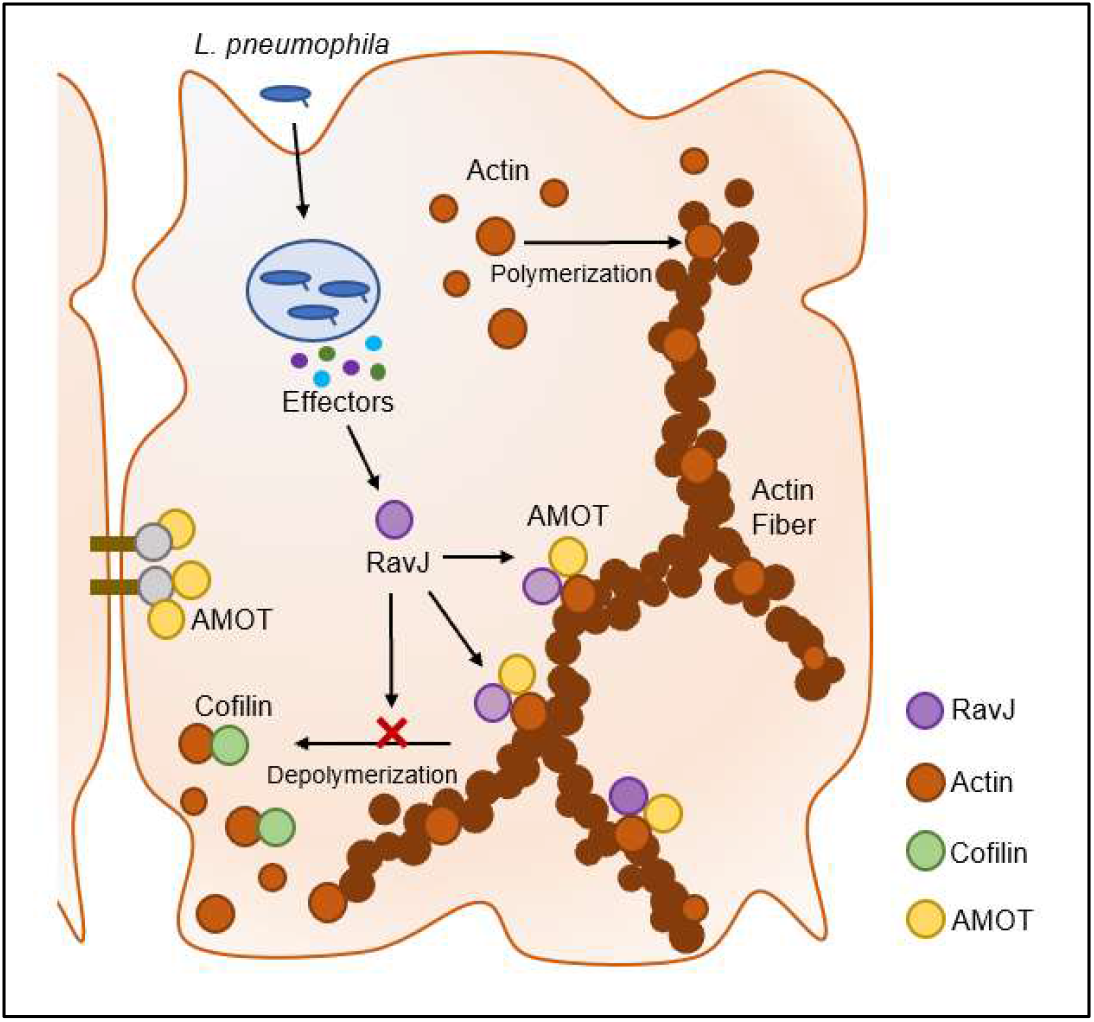
A predicted model of RavJ in actin cytoskeleton modulation. RavJ catalyzes the crosslink between actin and AMOT, which led to the blockage of the ADF/cofilin binding site in actin, resulting in stabilization of actin fibers in the cell cortex.

## Materials Availability Statement

All unique constructs and cell lines described in this article are available upon reasonable request from academic researchers. Please contact the corresponding author at for request of any materials.

## Acknowledgements

We thank Drs. Qing Deng, Matthew Olson and Chittaranjan Das (Purdue University) for helpful suggestions. We thank members of the Luo laboratory for helpful discussion. We also thank Dr. Uma K. Aryal from Purdue University and Dr. Jonathan Trinidad from the laboratory for Biological Mass Spectrometry at Indiana University for assistance in mass spectrometry analysis.

